# Coexisting morpho-biotypes unveil the regulatory bases of phenotypic plasticity in pancreatic ductal adenocarcinoma

**DOI:** 10.1101/2022.12.12.520054

**Authors:** Pierluigi Di Chiaro, Lucia Nacci, Stefania Brandini, Sara Polletti, Benedetta Donati, Francesco Gualdrini, Gianmaria Frige, Luca Mazzarella, Alessia Ciarrocchi, Alessandro Zerbi, Paola Spaggiari, Iros Barozzi, Giuseppe R. Diaferia, Gioacchino Natoli

## Abstract

Intratumor morphological heterogeneity predicts clinical outcomes of pancreatic ductal adenocarcinoma (PDAC). However, it is only partially understood at the molecular level and devoid of clinical actionability. In this study we set out to determine the gene regulatory networks and expression programs underpinning intra-tumor morphological variation in PDAC. To this aim, we identified and deconvoluted at single cell level the molecular profiles characteristic of morphologically distinguishable clusters of PDAC cells that coexisted in individual tumors. We identified three major morpho-biotypes that co-occurred in various proportions in most PDACs: a glandular biotype with classical epithelial ductal features; a biotype with abortive ductal structures and expressing a partial epithelial-to-mesenchymal transition program; and a poorly differentiated biotype showing partial neuronal lineage priming and absence of both ductal features and basement membrane. The identification of PDAC morpho-biotypes may help improve patient stratification and therapeutic schemes taking into account the spectrum of actionable targets expressed by coexisting tumor components.

## Introduction

Pancreatic ductal adenocarcinoma (PDAC) is the most lethal among common solid malignancies. While 5-year survival following surgery increased from 1.5% to 17% in the last 50 years, it showed only marginal improvements in the majority of PDAC patients (*ca*. 85%), who could not undergo surgery due to locally advanced or metastatic disease at diagnosis and whose median life expectancy is shorter than 6 months (Bengtsson et al., 2020).

Treatment failures in PDAC likely have several concurrent causes, one of them being the extensive cellular and architectural intra-tumor heterogeneity (Adsay et al., 2005; Del Poggetto et al., 2021; Hayashi et al., 2020; Klöppel et al., 2000; Nagtegaal et al., 2020). Within an abundant and dense fibrotic stroma accounting for up to 90% of the tumor mass, PDACs contain in variable proportions groups of tumor cells of different size, shape and subcellular features organized in growth patterns ranging from pseudo-glandular structures to compact nests (Verbeke, 2016). While such intratumor morphological heterogeneity is not part of standard pathological reporting, it has obvious prognostic impact (Verbeke, 2016), suggesting that the extent of morphological variation in PDAC is evidence of the complexity of the underlying tumor biology. For instance, a grading system that scores different and coexisting morphological patterns is a strong and independent predictor of prognosis (Adsay *et al*., 2005). Moreover, a recent pattern-based morphological classification system accurately predicts prognosis (Kalimuthu et al., 2020).

At the molecular level, PDAC intratumor heterogeneity is paralleled by the variability of gene expression programs revealed by single cell transcriptional analyses (Chan-Seng-Yue et al., 2020; Juiz et al., 2020; Raghavan et al., 2021), with high PDAC cell heterogeneity correlating with worse overall survival (Hwang et al., 2022). These data point to the notion that the coexistence of heterogeneous tumor cells lays the ground for swift adaptation to therapy or to therapy-induced selection of aggressive and treatment-resistant tumor cells. However, single cell RNA-seq data do not clarify the relationship between gene expression programs and variable morphological patterns, thus hindering a complete understanding of the molecular circuitry underlying morphological heterogeneity.

In the last decade, efforts to categorize PDACs mainly on the basis of bulk transcriptomic data led to several molecular classification systems. Most of them converge on the existence of a *classical* (or *pancreatic progenitor*) signature characterized by the preferential expression of typical endodermal or epithelial differentiation genes, and a series of partially overlapping, yet distinct signatures associated with loss of endodermal lineage fidelity, and referred to as *quasi-mesenchymal, basal* or *squamous* signature, with the different names reflecting the most over-represented gene expression program identified in the different studies (Bailey et al., 2016; Collisson et al., 2019; Collisson et al., 2011; Moffitt et al., 2015). While the *quasi*-mesenchymal signature includes genes involved in epithelial-to-mesenchymal transition (Nieto et al., 2016), a common occurrence in epithelial tumors, the basal and squamous signatures include regulators of basal cells, the stem cells of various glandular and stratified epithelia, and genes expressed by their differentiating progeny such as keratins characteristic of stratified epithelia. Critically, bulk tumor analyses average the transcriptomes from different coexisting tumor cells, with the classification of each tumor reflecting the most abundant population and with the contribution of less represented tumor components being instead systematically overlooked. These averaging effects in such a heterogenous tumor may explain why all these classification systems have overall limited clinical impact (Collisson *et al*., 2019) and are inferior to morphological classifications at predicting prognosis (Kalimuthu *et al*., 2020).

When considering mutational profiles, most PDACs display frequent genetic alterations in four main genes (*KRAS, TP53, CDKN2A, SMAD4*) in addition to less pervasive mutations or copy number alterations in genes encoding chromatin regulators and transcription factors (Hayashi et al., 2021). However, the contribution of genetic diversity to heterogeneity of PDAC cells, albeit detectable (Chan-Seng-Yue *et al*., 2020), appears to be lower than the contribution of instructive signals generated by the tumor microenvironment (TME) (Grunwald et al., 2021; Raghavan *et al*., 2021; Tu et al., 2021). In other words, different PDAC cell states may mainly be driven by local tissue signals rather than being hardwired in distinct cancer cell genomes. Indeed, while primary or stable cell models (organoids and cell lines) faithfully recapitulated PDAC genetic profiles, they failed to recapitulate their gene expression complexity; however, they could be instructed to adopt alternative states by altering cell culture conditions (Raghavan *et al*., 2021).

In order to fill the knowledge gap between intra-tumor morphological variation and molecular understanding of PDAC heterogeneity, in this study we aimed to determine the regulatory circuitries underlying the different morphological patterns that coexist in individual PDACs. We identified three main *morpho-biotypes*, defined as combined morphological patterns, molecular profiles and functional properties, that coexist in various proportions in most PDACs. Overall, our data indicate that PDACs are ensembles of morphologically discrete, spatially organized groups of tumors cells with distinct gene expression programs and functional properties, with direct implications for target-specific combinatorial therapies.

## Results

### Identification of coexisting morpho-biotypes in PDAC

We used laser micro-dissection (LMD) to isolate from primary PDACs of treatment-naïve patients (n=30, **Suppl. Table 1**) multiple morphologically distinguishable areas composed of up to *ca*. 200-500 cells each. Specifically, from each PDAC we isolated multiple areas selected on the basis of their different morphology, with each area being initially assigned to either of two broad categories: gland-forming or non-gland-forming tumor areas (Kalimuthu *et al*., 2020) (**Figure 1A**). To minimize contamination by non-tumor cells, we used immunohistochemical staining for pan-Cytokeratin on a consecutive section to guide the definition of the tumor cell borders. In addition, settings were optimized to keep laser cut width below 10μm, thus maximizing precision of the dissection.

**Figure 1.**
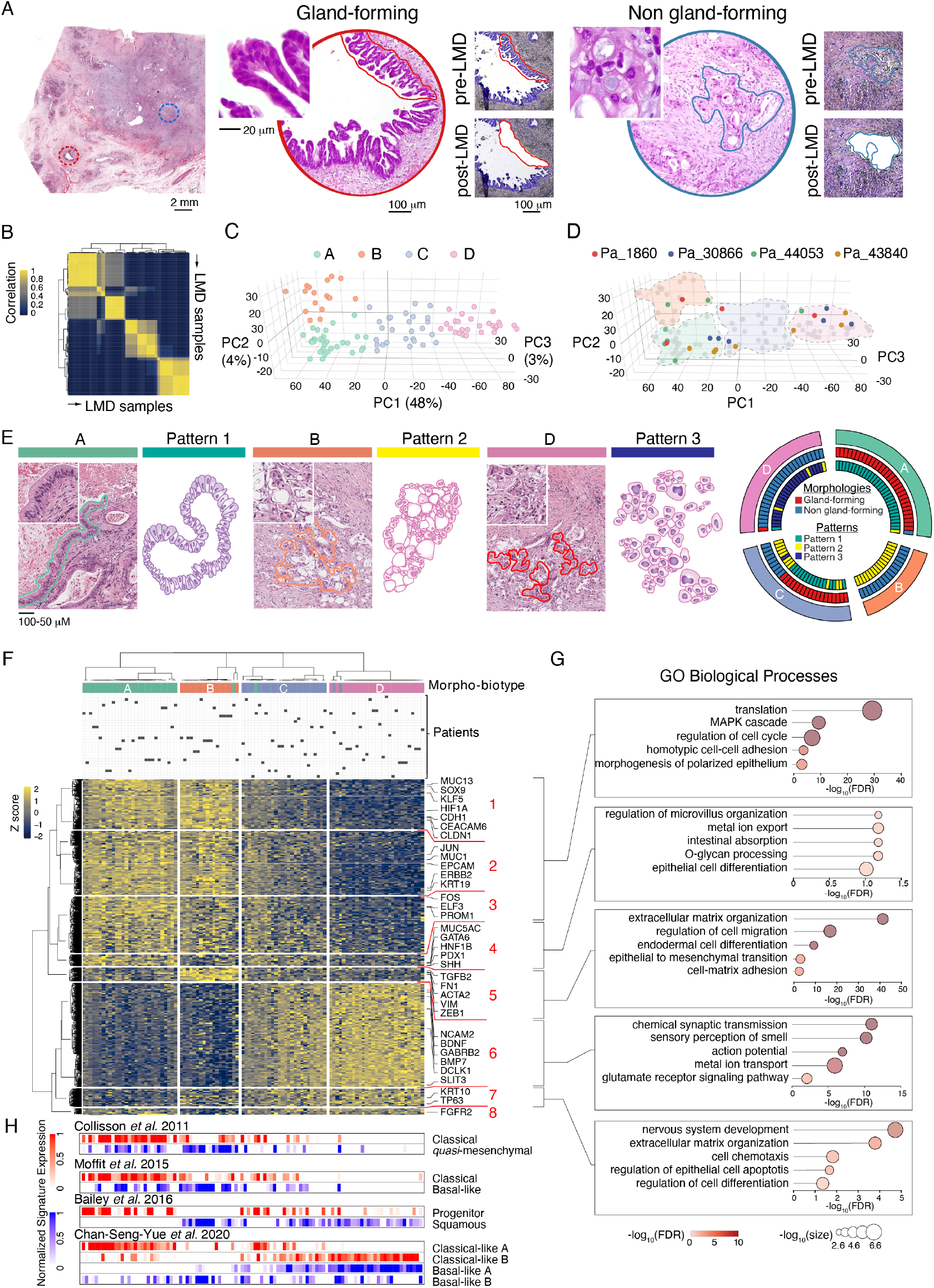
Three distinct morpho-biotypes coexisting in PDACs. **A**) Hematoxylin and eosin (H&E) images of a gland-forming (red circle) and a non-gland-forming area (blue circle) coexisting in the same PDAC. The same regions were matched on a consecutive section prepared for laser microdissection and imaged pre- (upper panels) and post-cut (lower images). Scale bars are indicated. **B**) Correlation matrix obtained by consensus clustering (k = 4, 1000 iterations) of the top 25% variable genes from the LMD-seq dataset (n=102 samples). **C**, **D**) 3D-PCA plots showing the distribution of the samples in the first three principal components. **C**) samples were colored by the four identified clusters (A, B, C and D) shown in panel **B**. **D**) Multiple samples from four different patients were colored to show their distribution in the four highlighted clusters (A: green, B: orange, C: blue and D: pink). **E**) Left: association of the microdissected areas (H&E images) to the morphological patterns derived from Adsay et al (Adsay *et al*., 2005) (schematic drawing depicting typical features). Right: circos plot of the identified sample clusters (A, B, C and D), the morphological components (gland forming or non-gland forming) derived from Kalimuthu et al (Kalimuthu *et al*., 2020) and the morphological patterns (patterns 1-3) derived from Adsay et al (Adsay *et al*., 2005). Scale bars: 100 mm; insets: 50 mm. **F**) Heatmap showing the four sample groups derived from differentially expressed genes clustered by graph-based clustering approach. Samples were re-clustered using ConsensusClusterPlus (Wilkerson and Hayes, 2010) with Pearson’s correlation distance, k-means clustering algorithm and 1000 iterations. Differentially expressed genes identified based on a fold change ≥ ± 2 in either direction and an FDR ≤ 0.01 were clustered using KNN and the Louvain graph clustering method. Biotype annotations are shown for each sample. **G**) Representative Gene Ontology classes (GO, biological processes) enriched in the indicated gene cluster. The dot color scale indicates the FDR while the dot size shows the frequency of the GO term in the GO Annotation (GOA) database. **H**) Gene expression scores in each sample using the indicated bulk gene signatures.

The microdissected PDAC areas (n=102; 1-7 per patient) were subjected to cDNA synthesis by random priming followed by RNA-seq (**Suppl. Table 2**), resulting in an average of 9,513 expressed genes per micro-dissected sample (range: 7,590-10,829). These numbers are in line with those obtained by standard RNA-seq while largely exceeding the number of transcripts detected per cell in single cell RNA-seq approaches. We applied consensus clustering (Monti et al., 2003) on the top variable genes followed by principal component analysis (PCA) to obtain an unsupervised classification of tumor areas (**Suppl. Table 3**). We identified four clusters with high intra-group correlations (**Figure 1B**) that were well separated by the first three principal components (**Figure 1C**).

While PC1 explained most of the variance (48%) observed between group A and B on the one hand, and group D on the other, PC2 separated group A from B (4% of the variance). Group C was intermediate between A and D. Importantly, tumor areas microdissected from the same patient clustered into different groups together with areas from other patients, suggesting that in spite of the high inter-patient variability characteristic of PDAC (Hwang *et al*., 2022), tumor areas with highly similar gene expression profiles can be detected across patients (**Figure 1D**).

Retrospective examination of histological data allowed us to assign different gene expression clusters to morphologically different areas that we could categorize into three main groups that broadly matched the morphological patterns previously reported to describe PDAC heterogeneity spectrum and to possess prognostic value when combined into a grading score (Adsay *et al*., 2005) (**Figure 1E**).

*Pattern 1 areas*, associated with the gene expression cluster A (**Figure 1E** and **Suppl. Fig. 1A**), consisted of epithelial duct-like structures in a wide gradation of sizes and architectures, from tubular and elongated glands to papillary and finger-like patterns. Tumor cells in these areas ranged from tall columnar cells resembling mucin-producing foveolar cells of the stomach, to cuboidal cells similar to those of the pancreato-biliary ductal system, indicating an overall extensive loss of lineage fidelity. In spite of this variety, most malignant cells displayed a clear cytoplasm related to the presence of abundant mucin or glycogen.

*Pattern 2 areas*, associated with the gene expression cluster B (**Figure 1E** and **Suppl. Fig. 1A**), included ill-defined glands with incomplete borders and irregular formation of multiple abortive lumina (*cribriform* architecture); these *cribriform units* displayed several small holes in between cells, which may represent rudimentary gland lumina, as well as frequent vacuolated tumor cells in which excess cytoplasmic mucin pushed the nucleus to the cell periphery (*signet-ring cells*).

*Pattern 3*, associated with the gene expression cluster D (**Figure 1E** and **Suppl. Fig. 1A**), consisted of *non-glandular, diffuse round cell clusters* in which cancer cells were arranged into sheets or nests with no architectural organization. Such clusters contained areas of (*i*) small packets of round neoplastic cells with variable compaction, intercellular bridges and minimal keratinization, (*ii*) large pleomorphic and anaplastic cells with prominent nuclear atypia and hyperchromatic nuclei and abundant eosinophilic cytoplasm; and (*iii*) nests of cells with solid or glassy-looking cytoplasm reminiscent of squamous cells with varying degrees of keratinization.

The gene expression cluster C showed a more complex association with both glandular and non-glandular areas and will be further discussed below.

Therefore, morphological diversity of small and distinct groups of tumor cells that coexisted in the same PDAC was associated with distinct gene expression programs. PDAC areas with distinct morphology and associated transcriptional programs are henceforth indicated as *morpho-biotypes* (or biotypes).

Differential gene expression analysis (fold change ≥ 2 and FDR ≤ 0.01) followed by unsupervised clustering partitioned the differentially expressed genes into eight clusters. This allowed us to highlight transcriptional differences and similarities among the morpho-biotypes (**Figure 1F** and **Suppl. Table 3**), which were then validated at single cell level (see below). The immunofluorescence staining of selected differentially expressed genes is shown in **Suppl. Fig. 1B**.

Specifically, morpho-biotypes A and B shared high-level expression of genes in clusters #1-3, which were enriched, among the others, with functional terms related to ribosome biogenesis and protein translation, suggesting high protein synthesis activity, as well as terms pertaining to cell-cell adhesion and epithelial polarization (**Figure 1G** and **Suppl. Table 4**). Notable genes in these clusters encoded: *i*) mucins (*e.g. MUC1; MUC13*); *ii*) CEACAMs (carcinoembryonic antigen-related cell adhesion molecules, *e.g. CEACAM6*); *iii*) epithelial adhesion molecules (*CDH1*, encoding E-cadherin; *EPCAM*, encoding the Epithelial Cell Adhesion Molecule; *CLDN1*, encoding the tight junction protein Claudin-1); *iv*) proteins involved in apical cell membrane organization (e.g. *PROM1*, encoding Prominin-1/CD133); and finally *v*) transcription factors associated with endodermal and/or pancreatic differentiation and also highly expressed in well-differentiated PDACs and cell lines such as SOX9, KLF5, ELF3 (Diaferia et al., 2016; Milan et al., 2021).

However, morpho-biotype A differed from B because of the selective expression of a group of genes (cluster #4) associated with endodermal differentiation and functions such as those encoding pancreatic lineage-determining transcription factors (GATA6, HNF1B and PDX1), as well as the *SHH* gene encoding Sonic Hedgehog, the main inducer of the desmoplastic reaction characteristic of PDAC (Bailey et al., 2008; Yauch et al., 2008). Conversely, biotype B selectively expressed genes (cluster #5) encoding proteins involved in extracellular matrix synthesis, cell migration and epithelial-to-mesenchymal transition (EMT) such as *TGFB2* (encoding the Transforming Growth Factor-beta2), *VIM* (encoding Vimentin), *FN1* (Fibronectin 1) and the transcriptional repressor ZEB1, a pivotal driver of EMT (De Craene and Berx, 2013).

Overall, biotype A (henceforth ***glandular*** morpho-biotype) was characterized by both a tubular architecture and the expression of the full spectrum of typical genes of the pancreato-biliary tree as well as endodermal lineage genes. Morpho-biotype B, while sharing with the glandular biotype the expression of an extended fraction of such genes, including many mucins, it showed on the one hand an incomplete glandular morphology with a dominant cribriform pattern, and on the other the downregulation of critical pancreatic lineage-determining transcription factors. Instead, genes involved in EMT as well as genes encoding extracellular matrix components (matrisome) were strongly upregulated in this biotype, a finding corroborated by single cell analyses (shown below). We tentatively named this morpho-biotype ***transitional*** to emphasize the induction of mesenchymal gene expression within groups of cells that however still retained the expression of a large share of ductal and endodermal genes, hinting at an incomplete or partial EMT as the regulatory basis for this specific morpho-biotype.

At the opposite end of the spectrum, morpho-biotype D was characterized by the extensive downregulation of most genes that define the glandular and transitional biotypes (clusters #1-5), and instead the upregulation of three large gene clusters (#6-8) encoding:

i. master regulators of clonogenicity and proliferation of stem and progenitor cells such as the transcription factor TP63, which is also overexpressed in various squamous cell carcinomas (Nylander et al., 2002) and in PDAC (Hayashi *et al*., 2020), and the protein kinase DCLK1, which marks cells with stem cell properties both in PDAC (Bailey et al., 2014) and in normal pancreas (Westphalen et al., 2016). Moreover, keratins of squamous epithelia such as *KRT10* (keratin 10) were selectively expressed in this biotype;
ii. *proteins associated with neuronal activities and functions*, such as *synaptic transmission and its regulation* (e.g., GABA receptors, NMDA-type glutamate receptor components, opioid receptors), *axonal migration* (e.g., SLIT-ROBO signaling complex components and neural adhesion molecules), *neurotrophic activity* (e.g., receptor tyrosine kinases including Neuregulin1 and NTRK family members), *voltage-gated ion channels* and *chemo-sensing* (e.g., olfactory receptor genes).

In particular, we noticed that a large fraction of genes (63/153) of the *Smell perception* gene expression program annotated in the Human Protein Atlas, was significantly enriched in the cluster #6 (FDR<0.0001, hypergeometric test) (**Figure 1G**, **Suppl. Fig. 2A-C**), suggesting a coordinated activation of such program rather than the stochastic activation of disparate neuronal genes. However, as it will become clear below, neuronal genes in this biotype are expressed at very low levels, a finding that is compatible with the priming of undifferentiated progenitors towards the neural lineage but not at all with a full deployment of a neuronal gene expression program. In addition, from a morphological viewpoint, these areas tended to coincide with the highly undifferentiated areas reported by pathologists (Verbeke, 2016). Therefore, based on combined morphological criteria and molecular data we deemed it appropriate to name this morpho-biotype as ***undifferentiated***, with neuronal gene expression representing a common, yet abortive lineage priming event in cancer cells uncapable to differentiate towards endodermal lineages. As mentioned above, the basal cell marker TP63 was associated to large and pleomorphic cell nests, a morphological feature commonly seen in undifferentiated carcinomas. Moreover, TP63-positive cells often co-expressed some neuronal enriched markers such as voltage-gated ion channels (KCNMB3 and KCNJ5) (**Suppl. Fig. 2D**).

An intriguing finding was the identification of tumor areas with a gene expression profile intermediate between the glandular and transitional biotypes on the one hand, and the undifferentiated biotype on the other (group C in **Figure 1E**). This *intermediate* or *hybrid* gene expression profile appeared to arise either from the low-level co-expression in individual cells of markers of different biotypes, a finding consistent with previous reports (Raghavan *et al*., 2021), or from the presence in a single ductal or non-ductal area of different cells expressing markers of alternative morpho-biotypes, with either sharp or gradual transitions between them (**Suppl. Fig. 1C**). Overall, this *hybrid* or *intermediate* profile may not represent a distinct entity but instead the consequence of the extreme loss of lineage fidelity commonly occurring in PDAC cells.

We next analyzed the relationship between the morpho-biotypes identified here and the PDAC molecular subtypes identified on the basis of bulk tumor, and thus averaged, gene expression signatures (Bailey *et al*., 2016; Collisson *et al*., 2019; Collisson *et al*., 2011; Moffitt *et al*., 2015).

The *classical* (or *progenitor-like*) signatures from several classification schemes were all consistently enriched in the *glandular* biotype and, as expected, also in the hybrid group (**Figure 1H**). An exception was represented by the *classical B* signature (Chan-Seng-Yue *et al*., 2020) that differently from the *classical A* was enriched in the undifferentiated biotype (**Figure 1H**). We notice that the classical A signature contains canonical pancreas differentiation and epithelial genes (such as *GATA6, GATA4, HNF4A, SHH, PROM1*, *CEACAM6, MUC1*); instead, while the classical B signature contains some endodermal genes (such as *FOXA2, MUC2* and *MUC13*), it is highly enriched for functional terms not immediately related to endoderm or pancreas (such as lipid synthesis and regulation of ion transport). The enrichment of this signature in the undifferentiated biotype was mainly due to a group of 36 genes that includes many brain-specific or brain-enriched genes (*e.g*., *GRIN2B, GRIP2, RPGRIP1, HECW2, ST6GALNAC3, LGI2, SCUBE1* etc).

Conversely, *mesenchymal* and *basal-like* gene signatures were enriched in the *transitional* biotype. Most importantly, the *undifferentiated* biotype was not significantly enriched for any of the signatures reported in Collisson et al. and in Moffit et al. (Collisson *et al*., 2011; Moffitt *et al*., 2015), which include only 62 and 50 genes, respectively. Instead, it was mildly enriched for the extensive (n=980 genes) *squamous* signature reported by Bailey et al. (Bailey *et al*., 2016). However, this squamous signature could not discriminate between the transitional and the undifferentiated biotypes as it was enriched in both (**Figure 1H**).

To validate the existence of the morpho-biotypes identified, we also used two orthogonal techniques. First, using the Nanostring GeoMx, we analyzed differences in gene expression programs associated with with gland- and non-gland-forming areas (n=8 patients among the 30 of the study; n=35 areas). With a predesigned panel of 1,834 genes, we could identify three clusters that were each specifically enriched for one the signatures characteristic of the three distinct biotypes, in addition to a cluster with a lower enrichment of all signatures and thus corresponding to the hybrid phenotype (**Suppl. Fig. 3A-D**). Second, we used multiplexed single-molecule FISH with probes corresponding to 32 selected genes highly expressed in the different biotypes. The results we obtained were consistent with the LMD-seq data and confirmed the existence of the three biotypes (**Suppl. Fig. 3E-H**).

### A Random forest-based classifier discriminating PDAC morpho-biotypes

We next used a machine learning approach in order to identify the minimal set of informative genes needed for a robust identification of morpho-biotypes. A Random Forest Classifier was trained on 50% (n=51) of the LMD-based gene expression dataset (training set, **Figure 2A**). To identify the smallest set of genes required for accurate classification, we applied recursive feature elimination (RFE) as a feature selection method to remove the weaker predictors, resulting in the identification of a 457 gene set. By allowing a 1% tolerance in model accuracy, this set could be reduced to 207 genes, indicating that the performance of the model is only marginally deteriorated by reducing the number of features used (**Figure 2B**). Predictor genes ranked by importance are shown in **Figure 2C**, while their levels of expression in the different morpho-biotypes are shown in **Suppl. Fig. 4A**. Notably, the identified predictors were significantly enriched for genes identified as differentially expressed among the three biotypes (OR=11.03, 95% CI=8.84,13.80; p-value < 0.0001 Fisher’s exact test) (**Figure 2C**). Finally, the 457 genes selected by RFE were used to cluster all the tumor areas. Unsupervised clustering showed that these genes were sufficient to recapitulate the initial clustering based on the expression of the top variable genes (**Figure 2D**), confirming the predictive power of the gene signature also in an unsupervised setting. Moreover, the resulting model enabled a precise classification (91% accuracy) of the remaining 50% (n=51) of the samples (testing set) (**Figure 2D**, **Suppl. Table 5**), indicating the robustness of the classifier.

**Figure 2.**
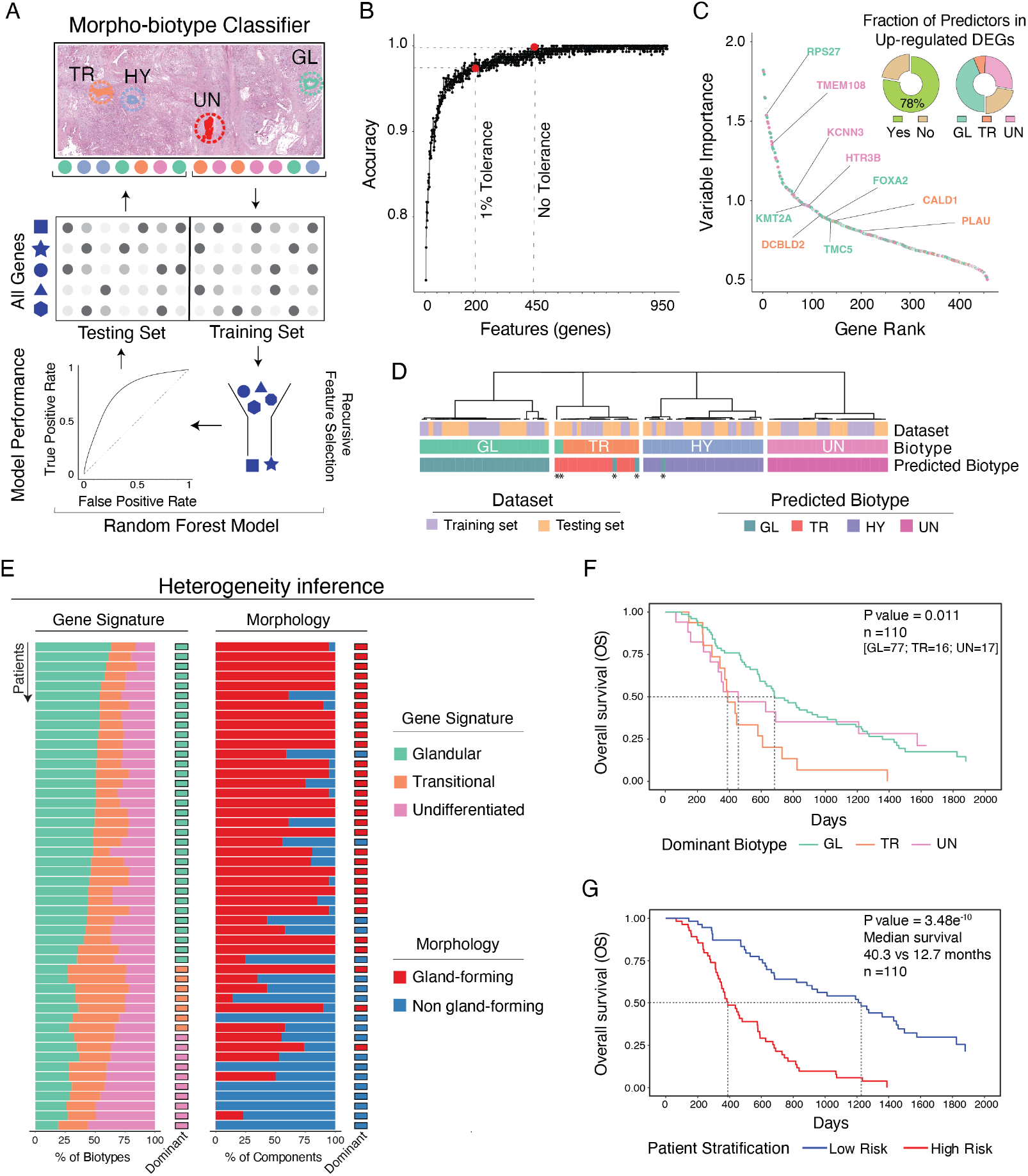
A Random Forest-based classifier discriminating PDAC biotypes. **A**) Schematic workflow of the Random Forest approach for the prediction of PDAC biotypes from LMD data. TR: transitional, HY: hybrid, UN: undifferentiated, GL: glandular biotype. **B**) Recursive feature elimination (RFE) analysis showing the optimal number of features (red dots and vertical black lines) at different percentage of tolerance in the model accuracy of the training set. **C**) Variable importance plot showing importance values (Gini index) of 457 genes selected by RFE and ranked in descending order from the most to the least important in the model. Each gene (dot) is colored according to biotype in which it was differentially expressed: A (glandular), green; B (transitional), orange; D (undifferentiated), pink; N/A, gray). Three selected genes for each biotype are highlighted. The donut charts show the fraction of the 457 genes matching all the up-regulated DEGs (left) or the up-regulated DEGs stratified into the three biotypes (right). **D**) Unsupervised clustering of all 102 LMD samples using ConsensusClusterPlus (Wilkerson and Hayes, 2010) with Pearson’s correlation distance, k-means clustering algorithm and 1000 iterations. Dataset (training and testing set) and the predicted biotype (GL: glandular, TR: transitional, HY: hybrid, UN: undifferentiated) annotations are shown for each sample. Asterisks indicate the five samples in the testing set that were not correctly predicted by the model. **E**) Left: the stacked bar plot on the left shows the inference of morpho-biotype composition of individual PDACs from bulk RNA-seq data, while the dominant biotype for each individual patient is shown on the right. GL: Glandular, TR: Transitional, UN: Undifferentiated. Right: stacked bar plot showing the percentage of gland-forming and non gland-forming morphological components as derived from Kalimuthu et al (Kalimuthu *et al*., 2020) in individual PDACs. The dominant morphological component for each individual patient is shown on the right of the panel. Only patients who died before the end of the study are shown (n=50). **F**) Kaplan-Meier plot of overall patient survival stratified by biotype (PACA-CA dataset, n=110). Log rank P-value is shown. Dashed lines indicate the median of survival for each biotype. **G**) Kaplan-Meier plot of overall patient survival for high-risk and low-risk groups (PACA-CA dataset, n=110). Log rank P-value is shown. Dashed lines indicate the median of survival for each group.

The random forest-based classifier was then applied to a different clinical dataset (Kalimuthu *et al*., 2020) to infer the morpho-biotype composition of individual PDACs based on bulk tumor RNA-seq data (**Figure 2E**). This analysis indicated on the one hand the presence of a dominant morpho-biotype and on the other the possibility to detect the co-occurrence of all the three morpho-biotypes in different proportions in different tumors. The inferred morpho-biotype composition was consistent with the reported morphological classification of the same cohort (n=66) (Kalimuthu *et al*., 2020). Overall, the model not only identified the dominant biotype but it could also be used to extrapolate the morpho-biotype composition of individual lesions, highlighting the co-existence of the three morpho-biotypes, and its correlation with the presence of different proportions of gland and non-gland forming tumor areas.

We next sought to determine the prognostic impact of the presence of different morpho-biotypes in individual PDACs. First, using the above classifier, we assigned each tumor of the same cohort ^11^ to one of three different classes based on the dominant morpho-biotype, and then determined the prognostic correlates of this assignment (**Figure 2F**). Tumors dominated by the transitional morpho-biotype showed significantly worse prognosis (HR=2.15, 95% CI=1.01,4.59; p-value < 0.05 Cox regression) than those dominated by a glandular morpho-biotype, a finding in line with the current literature (Collisson *et al*., 2019). Tumors with a dominant undifferentiated biotype showed a trend towards worse prognosis than those with a glandular dominance, although differences did not reach statistical significance (**Figure 2F**).

Survival analysis of patients showing different levels of expression of individual or combined morpho-biotype signatures showed that only the transitional signature and its combination with the undifferentiated one had significant, yet small prognostic impact (**Suppl. Fig. 4B**). We reasoned that since the three morpho-biotypes co-exist in individual PDACs, we may achieve better prognostic stratification by combining the expression of genes characteristic of different morpho-biotypes. We trained an elastic-net regression approach to extract a set of 23 prognostic genes from the 457 genes of the random forest classifier. The expression of these genes and their model coefficients, which approximately estimate the risk, were used to calculate a risk score for each patient and to stratify the cohort into a high-risk and a low-risk group. These two groups showed significantly different overall survival (p-value < 0.01 Log-rank test) (**Figure 2G**).

As an additional validation step, we used this 23 genes set to stratify patients from two additional independent cohorts. Although this patient stratification significantly impacted on the overall survival only in one of the two datasets (**Suppl. Fig. 4C**), the risk score was associated with worse prognosis in all cohorts analyzed (**Suppl. Fig. 4D**). Finally, even employing a more stringent four-fold cross validation in the initial cohort of the initial cohort (including the feature selection step to identify the prognostic genes; see **Methods**), confirmed the strong prognostic value of the risk groups (**Suppl. Fig. 4E**).

The prognostic genes derived above included genes belonging to all three morpho-biotypes but the level of expression of the individual genes of the glandular or undifferentiated biotype did not show a clear or a common association with patient stratification, since some of them were associated with better prognosis when higher expressed and some others when expressed at lower levels (**Suppl. Fig. 4F**). Notably, only one transitional biotype-specific gene (*FN1*, encoding Fibronectin), was strongly associated with poor survival when expressed at higher level.

Overall, the high prognostic power of combining genes associated with the three morpho-biotypes underscores the relevance of the co-existence of all the three morpho-biotypes for clinical assessment.

Finally, the prognostic model and its associated stratification strategy may help identify a high-risk group of patients who may benefit from target-specific combinatorial therapies since each morpho-biotype differentially expressed many targets of drugs currently in clinical use (**Suppl. Fig. 5** and **Suppl. Table 6**).

### Deconvolution of PDAC morpho-biotypes at single cell level

Although we micro-dissected very small and homogeneous tumor areas by accurately defining their borders, LMD-seq is nevertheless a population analysis in which different tumor and non-tumor cells may be co-isolated.

Therefore, we set out to use different single cell RNA-seq data-sets to determine the presence of the morpho-biotypes at the single cell level. First, we analyzed the enrichment of functional signatures extracted from single nucleus RNA-seq data (Hwang *et al*., 2022) in the morpho-biotypes (**Figure 3A-B**).

**Figure 3.**
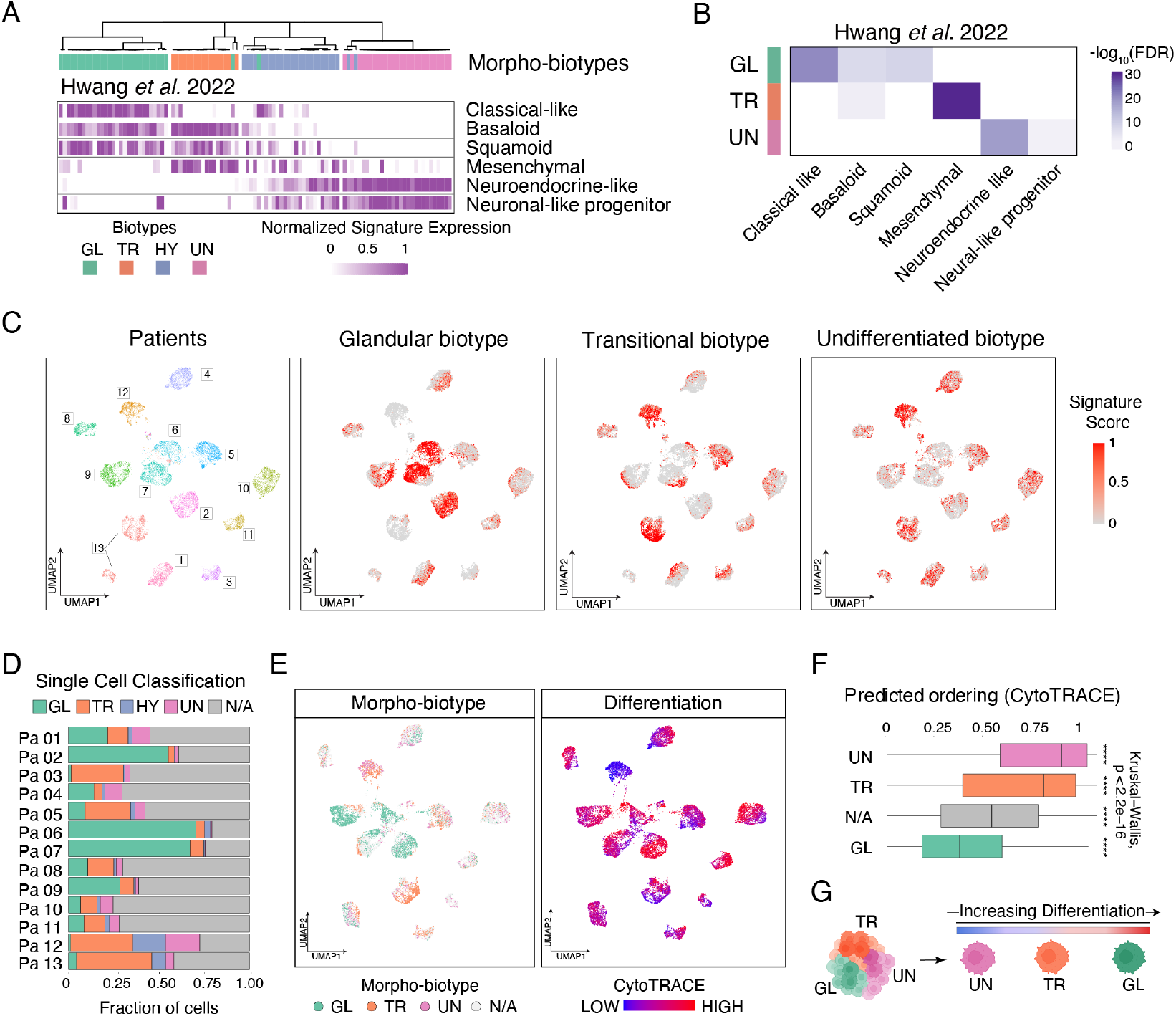
Deconvolution of PDAC biotypes at single cell level. **A**) Gene expression scores were calculated for each microdissected sample in the LMD dataset (clustered as in **Figure 1E**) using the indicated single cell gene signatures (Hwang *et al*., 2022). GL: glandular, TR: transitional, HY: hybrid, UN: undifferentiated. **B**) Heatmap showing the enrichment of the indicated tumor single cell transcriptional programs (Hwang *et al*., 2022) in each biotype-specific gene signature. The color scale indicates the statistical significance of the enrichment (−log_10_(FDR), hypergeometric test). GL: glandular, TR: transitional, UN: undifferentiated. **C**) UMAPs of malignant cells from 13 primary PDACs labeled by patient (left) or by gene expression score obtained for each biotype-specific gene signature (right). **D**) Stacked bar plot showing the fraction of single cells assigned to different biotypes based on gene expression scores (GL: Glandular, TR: Transitional, HY: Hybrid, UN: Undifferentiated, N/A: not assigned). **E**) UMAP of single malignant cells colored by biotype (left) or CytoTRACE (right) values (as proxies of differentiation). **F**) Box plot showing CytoTRACE values for the Glandular (GL, n=5498), Transitional (TR, n=3757) or Undifferentiated (UN, n=1286) biotypes and unclassified cells (N/A, n=12,000). Kruskal-Wallis P-value and twosided Wilcoxon rank-sum tests (group *vs*. rest) are shown. **G**) Schematic representation of differentiation state of PDAC cells of the three biotypes.

While a classical-like signature was enriched in the glandular morpho-biotype, we detected a selective enrichment in the transitional biotype of a mesenchymal signature that includes genes encoding components of the matrisome, the ensemble of hundreds of proteins constituting the extracellular matrix (discussed below). Notably, the *undifferentiated* biotype, showed a clear overlap with two neural-like (or neuro-endocrine) signatures that were mainly detectable in patients who underwent neo-adjuvant therapy (Hwang *et al*., 2022).

We next used a single cell RNA-seq data-set (Chan-Seng-Yue *et al*., 2020) to verify the enrichment of our morpho-biotype signatures in individual tumor cells identified on the basis of inferred copy number variants (Patel et al., 2014). The three morpho-biotype signatures showed complementary patterns of enrichment in cancer cells from all patients, confirming the co-existence of cells of the different biotypes in each tumor (**Figure 3C** and **Suppl. Fig. 6A**). We next assigned individual tumor cells from individual patients to the different biotypes. For each single cell, we first assessed the expression of all three morpho-biotype-specific signatures, and then assigned each cell to one or more biotype(s) based on the probability of observing by chance such expression values for each given biotype-specific signature (**Figure 3D**). While this analysis showed that the composition of each tumor was very heterogeneous, cells assigned to the undifferentiated biotype were a smaller fraction than those matching the glandular or the transitional ones. A large fraction of cells in each patient could not be assigned to any biotypes. This observation may simply relate to trivial threshold issues; however, it is consistent with previous studies reporting single cells that did not express programs corresponding to classical or basal signatures (Raghavan *et al*., 2021). Importantly, the level of expression of genes of the glandular and transitional biotypes signatures was consistently higher than that of genes of the undifferentiated biotype signature (**Suppl. Fig. 6B, C**). The relative rarity of undifferentiated biotype PDAC cells combined with the low level of expression of markers genes may contribute to explain why this gene signature has previously been overlooked in bulk transcriptome data but also in some scRNA-seq data set, in which genes expressed at low levels are inefficiently detected (Haque et al., 2017).

We next analyzed the differentiation state of PDAC biotype cells using CytoTRACE (Gulati et al., 2020), which uses the number of detectably expressed genes per cell as an indicator of their developmental potential, with undifferentiated cells expressing a high number of genes at low levels, and differentiated cells expressing at high level a comparatively lower number of differentiation-specific genes. We found that the glandular and the undifferentiated biotypes were at the opposite ends of the spectrum, being characterized by the highest and the lowest differentiation state, respectively, with the transitional biotype showing intermediate properties (**Figure 3E-F**). Therefore, based on this metric, cells of the undifferentiated biotype would resemble progenitor-like cells inclined to undergo neuronal lineage priming but also retaining the ability to sporadically differentiate towards alternative fates (such as metaplastic squamous differentiation) (**Figure 3G**). Importantly, the non-classifiable cells showed a differentiation state similar to the glandular cells, indicating that they may have been caught in a still incomplete, yet advanced differentiation stage.

### Differential basement membrane production in PDAC biotypes

One unexpected finding was the pattern of differential expression of genes related to the production of basement membranes, that in epithelial cancers represent structural barriers to tumor invasion and metastatization (Chang and Chaudhuri, 2019). Basement membranes are dense layers of extracellular matrix mainly composed of polymerized laminins and collagen IV, further crosslinked by additional proteins such as nidogen and perlecan and anchored *via* cell surface receptors to the underlying epithelial cells (Jayadev and Sherwood, 2017).

mRNA levels of basement membrane components were almost undetectable in the *undifferentiated* biotype, well expressed in the *glandular* biotype and, unexpectedly, further increased in the *transitional* biotype (**Figure 4A, B**), which is consistent with the enrichment in this biotype of both signatures related to extracellular matrix synthesis (**Figure 1G**) and the *mesenchymal* signature (**Figure 3A, B**).

**Figure 4.**
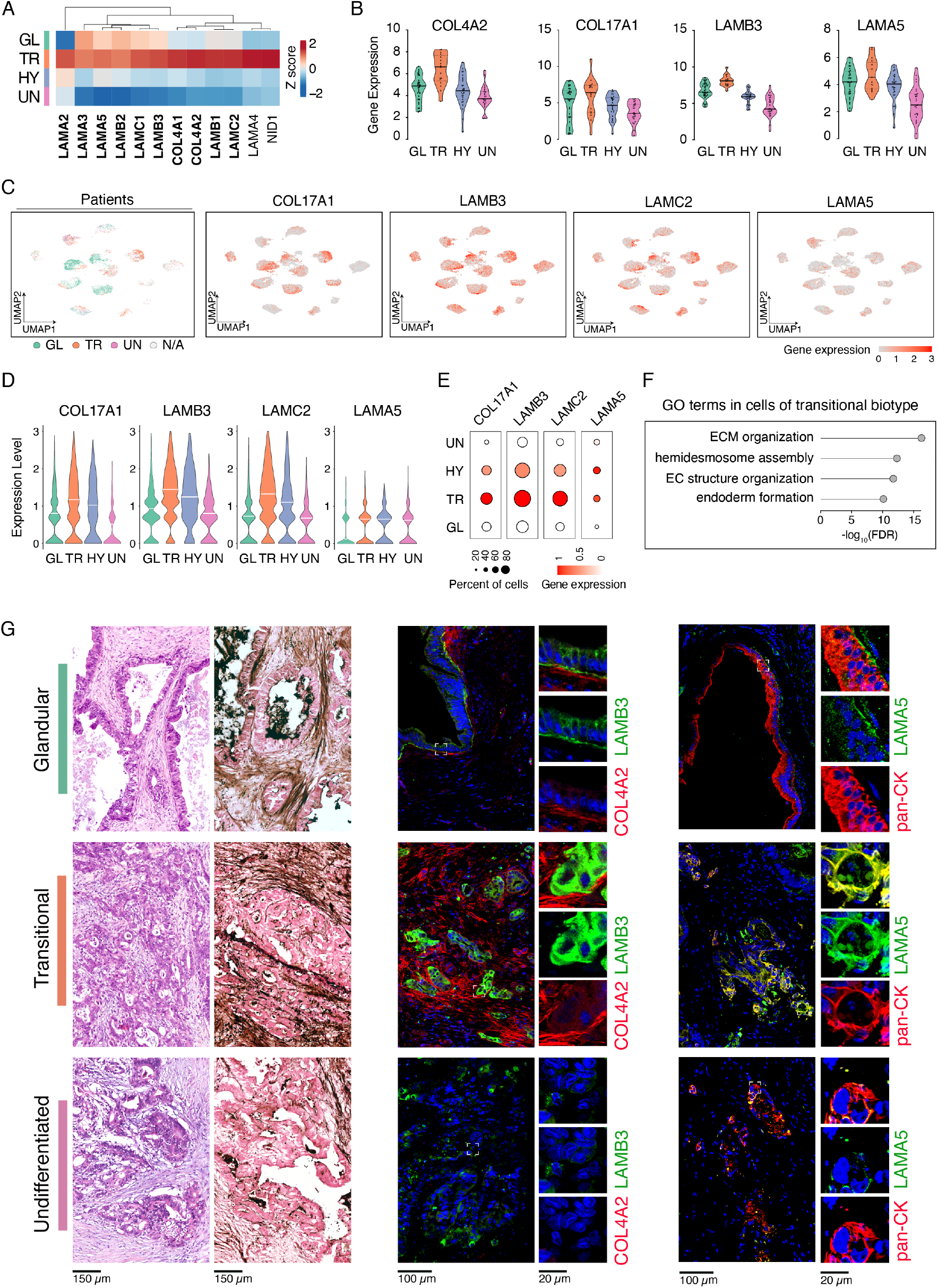
Production and assembly of basement membranes in PDAC biotypes. **A**) Heatmap showing the main components involved in the production of basement membranes (Jayadev and Sherwood, 2017) expressed in PDAC biotypes. Differential expressed genes are labeled in bold. **B**) Violin plots showing the expression levels stratified by biotype of representative differentially expressed genes validated at the protein level (panel **G**). Gene expression represents the log2(normalized read counts scaled by cDNA length + 1). **C**) UMAPs of single tumor cells from scRNA-seq data in 13 primary PDACs labeled by biotype (left); expression of selected genes encoding basement membrane components is shown (right). **D**) Violin plots showing the normalized gene expression of selected basement membrane components across the cells classified by biotype based on scRNA-seq data. **E**) Average gene expression level of selected basement membrane components in each biotype identified using scRNA-seq data. Dot color scale indicates the normalized mean expression while dot size shows the proportion of cells expressing in the biotype. **F**) Gene Ontology classes (GO, biological processes) enriched in the Transitional biotype cells identified in scRNA-seq data. **G**) H&E sections, PAS staining and IF images of the PDAC tumor areas showing the basement membranes in the different biotypes. Areas representative of the three biotypes in one PDAC patient (H&E and PAS-stained sections on the left) were immunostained for LAMB3 (green) and COL4A2 (red) in middle column panels and for LAMA5 (green) and KRT19 (red) in right column panels. Large immunofluorescent images showed the merged colors with DAPI-counterstained nuclei (blue). White boxes indicate the insert of area shown at higher magnification on the right, with red and green channels split in two images. Scale bars: H&E images 150 μm, PAS staining 150 μm, IF 100 μm, IF insert 20 μm.

The differential enrichment of these signatures may in principle be ascribed to contaminating fibroblasts isolated together with tumor cells upon microdissection. Therefore, we used scRNA-seq data sets to verify the differential expression of basement membrane components by malignant cells assigned to the different PDAC biotypes. The expression of several basement membrane components was significantly higher in cells of the transitional biotype compared to the other two, and specifically it was almost absent in malignant cells of the undifferentiated biotype (**Figure 4C-F**).

As a complementary approach, we used immunofluorescence to analyze the expression of Collagen IV, Laminin beta 3 (LAMB3), Laminin Alpha 5 (LAMA5), and periodic acid-Schiff (PAS) to stain basement membranes in PDAC tissue sections. Both *glandular* and *transitional* areas were associated with a strong staining of basement membrane components that were more abundant, but also more chaotically organized, in the transitional than in the glandular areas (**Figure 4G**). Notably, in the transitional areas, laminin B3 staining was mainly cytoplasmic rather than being deposited at the basal side of the cells, hinting at alterations in the control of basement membrane deposition. Finally, cell nests assigned to the *undifferentiated* biotype did not display any consistent or regular staining by PAS, suggesting the lack of a well-structured basement membrane (**Figure 4G**). Based on these data, it appears that differently from the other two morpho-biotypes, the undifferentiated one is not able to produce a basement membrane, with potential implications for containment of tumor cell invasion.

### Structural variation and mutational profiles in PDAC biotypes

Different biotypes may be generated as a consequence of alternative local microenvironmental signals acting on cancer cells (Raghavan *et al*., 2021; Tu *et al*., 2021) and/or be driven by the selection of clones characterized by distinct and characteristic genetic alterations.

We first used scRNA-seq data to infer copy number variations (CNV) and link them to cells assigned to different biotypes. An average biotype-specific CNV pattern was generated separately for each patient from the single-cell CNVs assigned to each biotype. Consistent with the high inter-patient transcriptional diversity, the inferred average CNV profiles clustered by patient rather than by biotype (**Figure 5A**). **Figure 5B** shows data from two representative patients.

**Figure 5.**
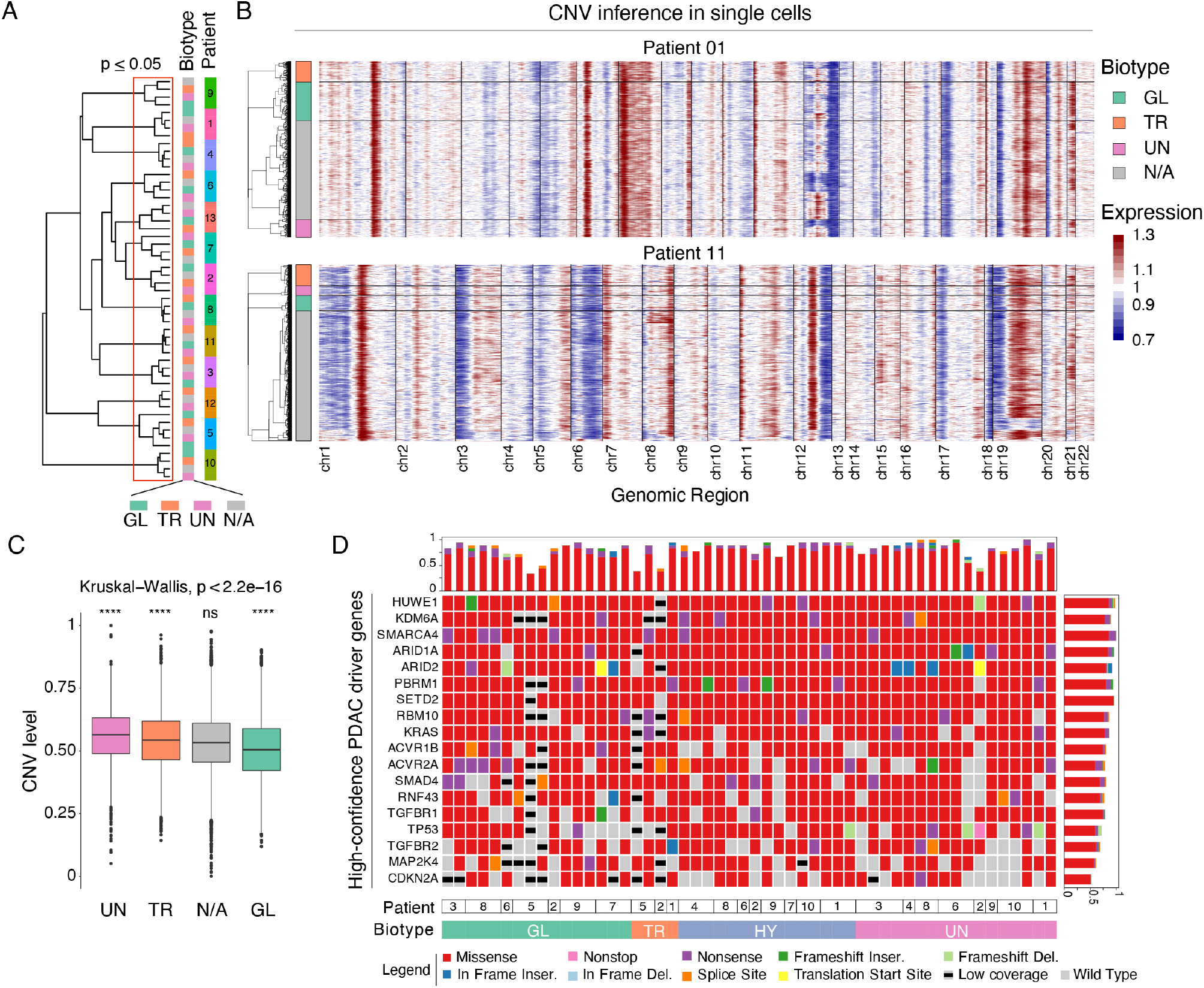
Genetic variation in PDAC biotypes. **A**) Hierarchical clustering of the mean of single-cell CNVs values (using Euclidean distance and the Ward.d2 method) for the Glandular (GL), Transitional (TR) and Undifferentiated (UN) biotypes and unclassified cells (not assigned, N/A) in each patient. The red rectangle indicates the cluster stability obtained by 1000 bootstraps (p ≤ 0.05). **B**) Heatmaps of inferred copy number variations (CNV) of single malignant cells from two representative patients clustered by biotype. Inferred amplifications are shown in red while inferred deletions are shown in blue. **C**) Box plot showing the distributions of CNV levels for the Glandular (GL, n=5498), Transitional (TR, n=3757) and Undifferentiated (UN, n=1286) biotype and unclassified cells (N/A, n=12,000). Kruskal-Wallis P-value and twosided Wilcoxon rank-sum tests (group versus rest) are shown. Biotypes are ordered from left to right based on decreasing median CNV level. **D**) Oncoprint showing non-synonymous somatic alterations of PDAC driver genes (Bailey *et al*., 2016) in 52 laser microdissected tumor areas obtained from 10 PDACs and grouped by biotype. Low coverage annotation indicates a low number of reads on more than half of all mutational sites called for a gene in a certain sample. Bar plots represent the fraction of somatic variants by gene (right) and by sample (top).

However, a higher number of CNVs was observed in cells of the undifferentiated biotype relative to the others and in particular to the glandular one, albeit differences were very small in magnitude (**Figure 5C**).

Next, we generated mutational profiles of 52 laser-microdissected areas obtained from 10 PDACs of our cohort and assigned to the different biotypes based on gene expression data (**Suppl. Table 7**). Each area was obtained from a section consecutive to the one used for RNA-seq. We used a custom amplicon-based panel covering 467 genes selected among cancer-predisposing (n=172) and driver or actionable genes (n=295, of which 62 in common with the hereditary ones) (Frige’ et al, manuscript in preparation) based on a revision of the literature updated to the PanCancer analysis (Bailey et al., 2018; Huang et al., 2018).

When considering both a restricted set of previously reported PDAC driver genes (Bailey *et al*., 2016) and the entire gene panel (**Figure 5D** and **Suppl. Table 7**), no significant differences in the mutational profiles were observed among morpho-biotypes. Overall, these data suggest that different biotypes may not arise because of the selection and expansion of clones with distinctively mutated driver genes.

### Transcriptional regulatory networks controlling biotype-specific programs

Finally, we set out to identify the transcriptional regulatory networks maintaining the different PDAC biotypes.

A large number of transcription factors (TFs) were differentially expressed in each biotype (**Figure 6A**), including known endodermal lineage TFs in the glandular biotype (*e.g*., SOX9, KLF5, HNF1B, GATA6, ELF3 and FOXA2) (Allen et al., 2011; Diaferia *et al*., 2016; Gao et al., 2008; Haumaitre et al., 2005; Martinelli et al., 2017; Milan et al., 2019), EMT-associated TFs in the transitional biotype (*e.g*., ZEB1 and GLIS2) (Balestrieri et al., 2018) as well as TFs associated with neural development (POU5F2, POU3F2, NOTO) but also stemness of basal epithelia (TP63) in the undifferentiated one.

**Figure 6.**
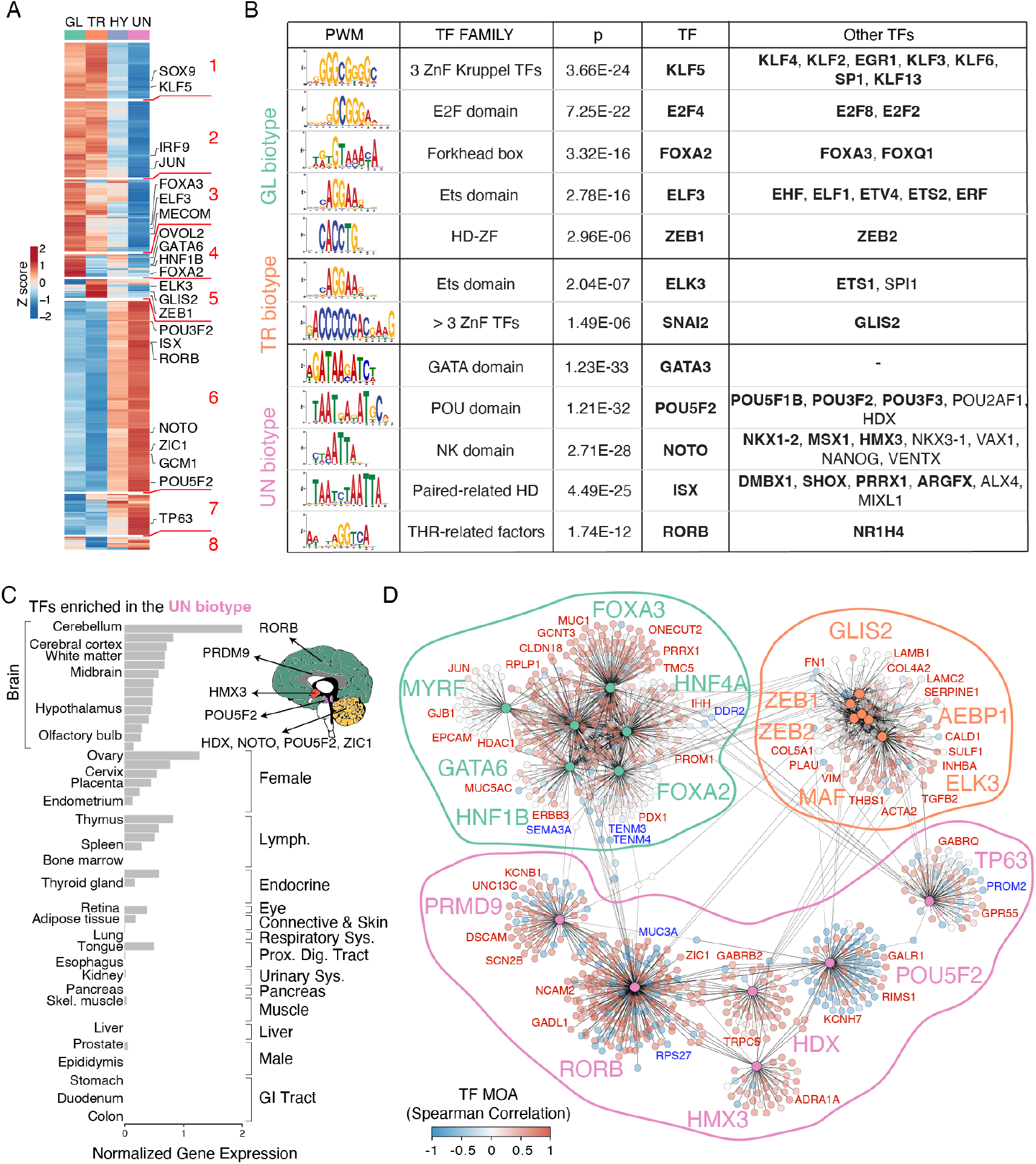
Biotype-specific transcriptional regulatory networks. **A**) Heatmap showing TFs differentially expressed in PDAC biotypes. Highly differential TFs (FC ≥ 3) and selected relevant TFs are shown. **B**) Motif enrichment analysis at the promoters of differentially expressed genes (from −500 bp to +50 bp relative to the annotated transcription start site). The table shows the PWMs overrepresented in one biotype relative to the others (p ≤ 1E-5), their TF family and their cognate TFs. TFs identified as differentially expressed in different biotypes are labeled in bold. **C**) Gene expression score for the TFs up-regulated in the Undifferentiated (UN) biotype in 55 tissues types from the consensus dataset of the Human Protein Atlas. Selected TFs enriched in brain and their tissue specificity are shown. **D**) Gene network representation of selected differentially activated TFs in the three biotypes as determined by ARACNe-VIPER. Each node represents a target gene that is predicted to be positively (red) or negatively (blue) regulated by its corresponding TF (TF MoA, mode of action of a TF based on Spearman correlation with its target genes).

To determine their possible involvement in enforcing biotype-specific regulatory networks, we first analyzed the statistical over-representation of TF DNA binding motifs in the promoters of differentially expressed genes characteristic of each biotype (**Figure 6B** and **Suppl. Table 8**). We identified motifs for endodermal lineage TFs such as KLF5, ELF3 and FOXA as enriched in the promoters of genes over-expressed in the glandular biotype. In addition, motifs bound by the EMT master regulator ZEB1 were also enriched in this group, consistent with ZEB1 acting as a transcriptional repressor (Balestrieri *et al*., 2018; Park et al., 2008). Binding motifs for the EMT activator SNAI2 (Nieto et al., 1994) were instead enriched in the transitional group. Consistent with gene expression data, promoters of genes over-expressed in the undifferentiated biotype were enriched for motifs bound by several neuronal lineage TFs such as POU5F2 and other family members and the homeobox TFs NOTO, NKX1-2 and HMX3 (**Figure 6B, C**).

Next, we sought to use our LMD-seq data to identify the transcriptional regulatory networks linking TFs and their target genes in the different biotypes. To this aim we first used a reverse engineering approach (Lachmann et al., 2016; Margolin et al., 2006) to reconstruct the regulatory networks from LMD data. We then used the VIPER algorithm (virtual inference of protein activity by enriched regulon analysis) (Alvarez et al., 2016) to determine the most active transcription factors based on the enrichment of their predicted target genes in each biotype (**Figure 6D** and **Suppl. Table 8**). In addition to linking TFs associated to endodermal lineage or EMT programs with their activated and repressed targets, this analysis confirmed the presence of an active neuronal-like network in the undifferentiated biotype, with possible direct links between neuron-enriched TFs and genes associated with functional categories characteristic of the nervous system.

## Discussion

PDACs are highly complex three-dimensional ductal systems embedded in an abundant stroma in which cancer cells in different regions are exposed to diverse microenvironmental signals and undergo further clonal diversification and selection, thus generating extensive intra-tumor heterogeneity also apparent at histopathological examination.

The objective of this study was to fill a major knowledge gap, namely the intersection between the heterogeneous morphological patterns that have long been known by pathologists not only to coexist in most PDACs but also to possess prognostic value (Adsay *et al*., 2005; Kalimuthu *et al*., 2020; Verbeke, 2016), and the distinct gene expression programs and genomic alteration profiles underlying PDAC heterogeneity. Thus, our approach differs from, and complements, on the one hand bulk tumor molecular classification schemes (which average transcriptional programs or mutational profiles associated with coexisting but distinct tumor cells), and on the other, single cell omics approaches that, while describing the full scope of tumor cell heterogeneity, fall short of precisely linking the observed molecular profiles to the extensive intratumor morphological variation characteristic of PDAC.

In this context, the previously described association of the classical and basal molecular signatures to gland and non-gland forming area, respectively (Kalimuthu *et al*., 2020), falls short of linking PDAC transcriptional complexity -as observed in single cell RNA-seq data or in our LMD-seq analysis-to the wide range of different morphologies observed in this cancer.

Based on our data, we introduced the concept of PDAC *morpho-biotypes* to highlight the notion that PDAC cells with different gene expression programs tend to be organized into morphologically distinct, identifiable and spatially discrete tumor areas (**Table 1**), which in principle may reflect two non-mutually exclusive scenarios: alternative tumor cell states instructed by different microenvironmental signals and/or localized selection and expansion of newly emerged clones. Analysis of the mutational profiles and chromosomal alterations in the different biotypes suggest a complex scenario whereby cells of the undifferentiated biotype tend to display a slightly higher burden of large-scale chromosomal alterations compared to the more differentiated cells of the glandular and transitional biotypes. While this observation may intuitively link poor differentiation capacity to corrupted genomic organization, the observed differences were of small magnitude and thus of unclear impact and fully compatible with a relevant role of microenvironmental signals either in preventing efficient differentiation or in actively promoting the maintenance of an undifferentiated state.

**Table 1.**
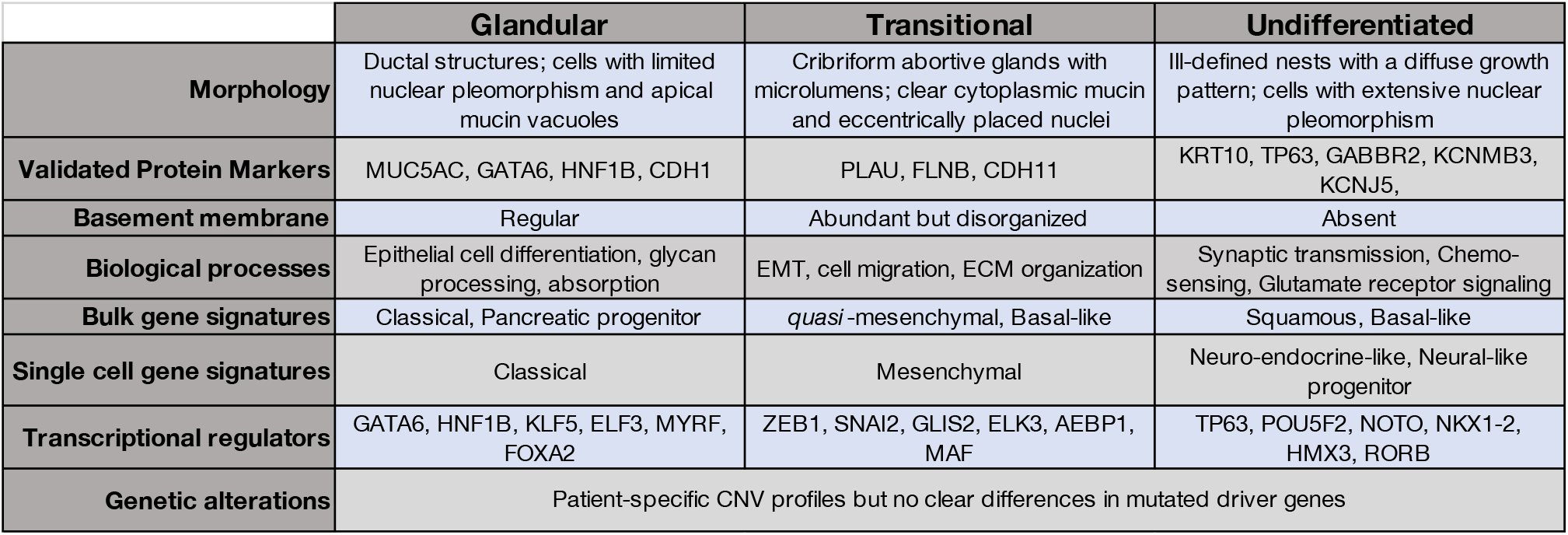
Summary of PDAC morpho-biotypes and their main features.

A most unexpected finding of this study was the identification of a low-level expression of a large number of neuronal genes by morphologically undifferentiated cells without any pseudo-glandular organization, and completely lacking the ability to produce basement membrane components. Because of the ectodermal origin of neurons, the identification of a neuronal-like gene expression profile in a tumor of an endodermal organ is surprising. However, the endodermal (and specifically pancreatic) origin of a subset of PDX1-positive neural progenitors contributing to the enteric nervous system has been reported in humans and mice (Brokhman et al., 2019; Seaberg et al., 2004; Smukler et al., 2011) and the direct development of neurons from endoderm was also shown in sea urchin (Wei et al., 2011), suggesting an evolutionary conserved developmental trajectory that may be pathologically activated in tumors. Indeed, neurogenic programs similar to those we retrieved have previously been associated with prostate cancer cell stemness (Zhang et al., 2016). Moreover, by mining published data we found neurogenic programs to be upregulated in primary and metastatic murine PDAC upon deletion of mesenchyme program-driving transcription factors (Carstens et al., 2021).

It is critical to stress, however, that while a large number of neuronal genes were specifically overexpressed in this biotype, their level of expression was low and thus indicative of only an initial priming towards the neuronal lineage. This interpretation is also consistent with the limited differentiation of these cells indicated by CytoTRACE. Moreover, the fact that the undifferentiated biotype also overlaps with basal and squamous signature genes is in principle compatible with the presence of undifferentiated cancer cells frequently undergoing priming towards the neuronal lineage but also capable of stochastically initiate alternative fates such as squamous differentiation, that in some cases reaches terminal stages. Indeed, cells of this biotype frequently expressed TP63, which maintains stemness and proliferative capacity in basal epithelial stem cells both in squamous and glandular epithelia. Notably, TP63^high^ cells have a characteristically high plasticity and are prone to multilineage priming (Claudinot et al., 2020).

Low levels of expression of this program in a rather small proportion of tumor cells may also readily explain why this gene expression profile was overlooked in bulk analyses, in which it was probably averaged out by more abundant cells expressing at high levels genes characteristic of more differentiated states. This signature was also overlooked in several single cell analyses, possibly due to the limited ability of scRNA-seq to robustly measure low-expressed genes. However, many neuronal genes were in fact included in the Classical B bulk signature (Chan-Seng-Yue *et al*., 2020) and a recent single nucleus RNA-seq data set (Hwang et al., 2020; Hwang *et al*., 2022) retrieved a similar signature which was enriched in patients who underwent surgery after neoadjuvant therapy, hinting at the possibility that these poorly differentiated cells may be positively selected in response to therapy. However, understanding the relevance of this undifferentiated biotype and specifically the relevance (if any) of the upregulation of a neuronal-like program by a subset of cancer cells will require extensive additional work in suitable experimental systems. Interestingly, proneural gene expression programs induced by palmitic acid treatment were associated with a high metastatic potential in oral squamous cell carcinoma (Pascual et al., 2021) and related to the promotion of perineurial invasion, which also represents a major dissemination pathway and an independent predictor of poor prognosis in PDAC (Schorn et al., 2017). However, we didn’t find any association between neuronal-like gene expression programs and perineurial invasion in PDAC (Nacci et al., unpublished data).

Because of the approach we used to isolate tumor cells, the morpho-biotypes reported here must be considered as broad categories in which both morphological and molecular heterogeneity exists. For instance, from a morphological point of view, within the glandular biotype a large spectrum of different epithelia can be observed, ranging from tall mucin-filled cells resembling gastric foveolar cells to villi-containing cells resembling the enterocytes, and finally cuboidal cells similar to the epithelial lining of the pancreato-biliary system. A similar degree of heterogeneity occurs in the diffuse, non-glandular areas characteristic of the undifferentiated biotype. From a molecular viewpoint, each biotype can include malignant cells displaying different combinations of the many PDAC gene expression programs identified in single cell RNA-seq data (Hwang *et al*., 2020; Hwang *et al*., 2022). Such heterogeneous tumor cells may represent either different functional states (e.g., cells activated by cytokines released by tumor-infiltrating immune cells) or cells intercepted at distinct developmental stages within an abnormally plastic and chaotic differentiation process. In this regard, the observation that basal, squamous and neuronal genes often tend to be co-expressed within the same tumor regions, points at the possible existence of pathological developmental niches in which stem or progenitor cells (such as basal-like cells) unable to generate fully differentiated ducts may instead be primed towards alternative differentiation programs such as the squamous-like and the neural-like one. Notably, a well-known finding in PDAC is the existence in undifferentiated tumor areas of cells that may belong to different lineages such as osteoclast-like giant cells and spindle cells resembling sarcomas (Demetter et al., 2021; Verbeke, 2016). Finally, we would like to highlight that while single cell RNA-seq permits to maximally deconvolute tumor cell heterogeneity, our approach instead enables the identification of groups of cells that because of their spatial proximity are likely linked by a progenitor-progeny relationship.

How will these data help improve the clinical approach to PDAC patients? First, these results suggest the possibility to implement novel PDAC grading schemes based on the representation of different biotypes in patient samples. Currently, PDAC grading has limited clinical relevance as it does not impact therapy and only some schemes have prognostic correlates (Milan *et al*., 2021). In the future, by expanding the data set reported in this study and by combining it with a digital annotation of the associated histopathological features, it may be possible to achieve a detailed description of biotype composition in each patient. Assuming that it will be possible to design combinatorial therapies based on tumor biotype composition, this novel grading approach may eventually contribute to inform therapeutic decisions. To achieve this objective, it will be critical to generate tractable experimental models recapitulating the properties of individual PDAC biotypes.

## Acknowledgements

We thank Andrea Viale (MD Anderson Cancer Center) and Silvia Monticelli (IRB, Bellinzona) for critical comments on the manuscript. This study was supported by AIRC, the Italian Association for Research on Cancer (AIRC Investigator Grant 20251 to G.N. and AIRC 5×1000 Grant ISM) and by the Italian Ministry of Health (grant GR-2016-02361721 to G.R.D). This work was also partially supported by the Italian Ministry of Health with the “Ricerca Corrente” and “5×1000” funds to the IEO IRCCS. P.D.C. is supported by an AIRC fellowship. The single cell RNA-seq datasets from PDAC samples used in this research were provided by the Ontario Institute for Cancer Research through funding provided by the Government of Ontario.

## Author contributions

Experimental design and conceptualization: PDC, LN, GRD, GN.

Experimental work and data generation: LN, PDC, SB, GF, GRD, SP, BD.

Data analysis: PDC, FG.

Clinical samples and pathology: AZ, PS.

Supervision: LM, AC, IB, GRD, GN.

Funding acquisition: GRD, GN.

Manuscript writing: PDC, LN, GRD, GN.

## Methods

### Patient cohort

Human PDAC specimens were provided by the Humanitas Research Hospital IRCCS (Rozzano, Italy) with written consent for tissue donation and under a protocol approved by the institutional ethical committee. All patients had a premortem diagnosis of PDAC based on histopathological analysis on the resected material and they did not receive any neo-adjuvant therapy before surgery. A summary of the patient cohort data is provided in **Supplementary Table 1**.

### Histological review

Hematoxylin and Eosin (H&E)-stained slides cut from formalin-fixed paraffin-embedded (FFPE) sections were reviewed by a gastrointestinal pathologist (P.S.) on the basis of biomarker studies and cytological appearance. H&E slides were scanned using an Aperio ScanScope XTsystem (Leica Biosystems) at 20x magnification and morphological distinct areas were drawn on the digital sections. Annotated areas were matched and aligned to the consecutive sections mounted on specific slide for microdissection (see below).

### Laser microdissection (LMD)

4 μm thick FFPE sections were mounted on polyethylene naphthalate membrane (PEN) slides (Leica cat. 11600289) previously UV-photoactivated in an UV crosslinker for 30 min (BLX-254, Bio-Link). FFPE sections were deparaffinized with two changes of xylene and then partially rehydrated in an ethanol gradient up to 75% EtOH.

Sections were then counterstained for 30 s with freshly prepared Cresyl Violet (0.8% Cresyl Violet in 60% EtOH and 4 mM Tris–HCl, pH 8.0), washed twice in 75% EtOH and air dried completely before proceeding to the microdissection. Single or multiple areas were drawn to collect 200-500 cells per sample using a UV-based LMD7 laser microdissection system (Leica Microsystems) at 20x magnification. Whenever possible, multiple samples of morphologically distinct are were collected in each patient. Microdissected areas were collected by gravity into strip-tube caps and immediately processed for RNA or DNA library preparation.

### RNA-seq library preparation

Microdissected areas were digested with 10 mU of Proteinase K (ThermoFisher, #AM2546) in 6 ml of digestion buffer (0.05% Triton X-100, 30 mM Tris-HCl pH 8.0) for 1 hour at 60C. 1 ml of 5 mM Proteinase K inhibitor (Calbiochem, #539470) was added to stop the reaction and samples were stored at −80 C until library preparation.

Samples were then processed using the SMART-seq Stranded kit (Cat. 634444, protocol version 051018; Takara Bio USA, Inc.) according to the manufacturer’s protocol with minor modifications. Briefly, the fragmentation step was omitted and RNA molecules were copied into first strand cDNA by reverse transcription using random primers and a template-switching oligo (TSO). Sequencing adapters and indexes were added to single-stranded cDNA with 5 cycles of PCR. cDNA originating from ribosomal RNA was depleted using scZapR in the presence of the mammalian-specific scR-Probes following manual instructions. The resulting ribo-depleted library fragments were amplified with 14 cycles of PCR, purified with AMPure beads and profiled for size distribution on an Agilent 2200 TapeStation with High Sensitivity D5000 reagent kits.

Libraries were quantified by QuantiFluor assay run on a Glomax Explorer instrument (Promega) and sequenced on an Illumina NovaSeq 6000 platform to obtain 50 bp paired-end reads.

### Gene panel mutational analysis

Areas selected for microdissection corresponded to sections consecutive to those used for RNA-seq. Microdissected areas were digested with 12 ml of Direct Reagent (Ion AmpliSeq™ Direct FFPE DNA Kit, MAN0014881, ThermoFisher Scientific) for 30 min at 65 C. The digested samples were subjected to DNA target enrichment by 15 cycles of PCR amplification using the Ion AmpliSeq Library Kit Plus (MAN0017003, ThermoFisher Scientific) and a custom pool of primers (GerSom). GerSom is a custom amplicon-based panel covering 467 genes selected among cancer-predisposing and driver genes, for a total sequencing space of 2.2 Mb. Panel design details are being separately submitted (Frigè et al., in preparation) and are available upon request. After partial amplicon digestion and phosphorylation, the fragments were purified and used for adapter ligation and library amplification using the NEBNext Ultra II Fs DNA Library kit for Illumina (NEB # E7805, New England BioLabs) according to the manufacturer’s instructions. Quality-checked and quantified libraries (see above) were pooled and sequenced (150 bp paired-end reads) on an Illumina NovaSeq 6000 platform.

### Immunofluorescence

FFPE sections were deparaffinized, rehydrated and subjected to heat-induced antigen retrieval in EDTA buffer (1 mM EDTA pH 8.0, 0.05% Tween-20) using pressure cooker (TintoRetriever, BioSB). After blocking, sections were incubated overnight at 4 C with the following primary antibodies: anti-pan-Cytokeratin Alexa Fluor^®^ 488 conjugated (2.5 μg/ml, Novusbio #NBP2-33200AF488), anti-KRT10 (1:100, Biocare Medical, cytokeratin HMN #CM127A,C), anti-FLNB (0.5 μg/ml, Sigma-Aldrich, #HPA004747), anti-TP63 (1:75, Dako, #M7317), anti-HNF1B (0.4 μg/ml, Sigma-Aldrich, #HPA002083), anti-MUC5AC (1:50, Dako, #M7316), anti-GATA6 (10 μg/ml, Santa Cruz, #sc-9055X), anti-CDH1 (1:50 mg/ml, Dako, #M3612), anti-CDH11 (10 μg/ml, ThermoFisher Scientific, #32-1700), anti-CEACAM6 (2 μg/ml, Abcam, #ab137859), anti-GABBR2 (1 μg/ml, Sigma-Aldrich, #HPA031684), anti-PLAU (0.5 μg/ml, Sigma-Aldrich, #HPA008719), anti-LAMB3 (10 μg/ml, Sigma-Aldrich, #AMAb91160), anti-Collagen IV alpha 2 (3 μg/ml, Atlas Antibodies, #HPA069337), anti-LAMA5 (5 μg/ml, Sigma-Aldrich, #AMAb91124), anti-KCNMB3 (1 μg/ml, Sigma-Aldrich, #HPA015665), and anti-KCNJ5 (0.5 μg/ml, Sigma-Aldrich, #HPA017353).

After washing, anti-rabbit Rhodamine-Red-X conjugated and anti-mouse Alexa Fluor^®^ 647 conjugated fluorophore-labeled F(ab)2 donkey secondary antibodies were used (Jackson Immunoresearch). Sections were DAPI counterstained and 3×3 Z-stacked large images (9-5 mm depth, 11-7 stacks) were acquired using a Yokogawa Spinning Disk Field Scanning Confocal System (CSU-W1, Nikon Europe BV, Amsterdam, Netherlands) equipped with 405, 488, 561, 640, 785 nm lines of solid-state lasers, 40x/1.15NA water immersion objective lens and a Prime BSI sCMOS camera (Teledyne Photometrics, Tucson, AZ).

### Jones Stain (methenamine silver-Periodic acid-Schiff stain)

The Jones Stain Kit (Abcam, #ab245883) was used according to the manufacturer’s protocol. Images were acquired using Aperio ScanScope XTsystem (Leica Biosystems) at 20x magnification.

### GeoMx Digital Spatial Profiling (DSP)

5 mm glass-mounted FFPE sections were baked at 65C for 1 h and manually prepared using the manufacturer supplied V2.0 protocol (MAN-10115-01). Slides were hybridized to UV-photocleavable barcode-conjugated RNA probe set (Cancer Transcriptome Atlas Panel) overnight at 37C using a Hyb EZ II hybridization oven (Advanced Cell Diagnostics). After stringent washes with 2X SSC and formamide at 37 C, slides were stained with fluorescently labeled morphology markers (Pan-cytokeratin, CD68 and Syto13) for 1 hour to distinguish tumor cells from the tumor microenvironment. Slides were loaded onto the GeoMx DSP instrument for imaging, tumor area selection, barcode cleavage and region of interest (ROI) collection (3-4 ROIs per PDAC patient). Library preparation was performed according to manufacturer instructions (Nanostring DSP-Genomics Library Preparation Protocol MAN-10117-01). Pooled libraries were sequenced on an Illumina NextSeq2000 platform (75 bp paired-end reads).

### Multiplexed RNA smFISH

5 mm glass-mounted FFPE sections were baked at 65C for 1 hour, deparaffinized, rehydrated and subjected to heat-induced antigen retrieval in TE buffer (10mM Tris-HCl, 1mM EDTA, pH 8.0) using a steamer at 98 C for 20 min. After blocking in 5%BSA, a rabbit polyclonal antibody against KRT19 (10ug/ml, Novusbio # NBP1-78278) was incubated for 1h at room temperature and revealed with a Pacific Orange-conjugated secondary antibody (1:50, ThermoFisher, # P31584). After DAPI counterstaining, four representative heterogeneous tumor areas were acquired as Z-stacks using a Yokogawa Spinning Disk Field Scanning Confocal System (CSU-W1, Nikon Europe BV, Amsterdam, Netherlands) equipped with 405, 488, 561, 640, 785 nm lines of solid-state lasers and 60x/1.4NA oil immersion objective lens and a Prime BSI sCMOS camera (Teledyne photometrics, Tucson, AZ). After washing in PBS, samples were subjected to 8 rounds of hybridization with 4 probe sets at 4nM each according to the manufacturer’s protocol (Molecular Instruments). After overnight hybridization and washes, probes were amplified using a mixture of metastable paired hairpins (HCR amplifiers) (Choi et al., 2018) conjugated with Alexa488, Alexa594, Alexa647 and Cy7 fluorophores at 60 nM each in Amplification Buffer (Molecular Instruments) for 45 min at room temperature. After washing, background autofluorescence quenching (VectorLabs, #SP-8400) and DAPI counterstaining the same tumor areas were imaged as before.

DNA probes were digested with 0.5U/ml RNase-free DNase I (Roche, #04716728001) for 1h at RT to perform a multi-day hybridization routine. After probe displacement, samples were first washed several times in Wash Buffer and then hybridized with a new probe set as described above.

For quantification of the RNA transcripts, the images acquired after each round of hybridization were processed using starfish (Perkel, 2019). To correct for microscope stage drift during the multiple rounds of imaging, raw images were first registered by taking a maximum intensity projection along the z direction in each channel and then were aligned using a sub-pixel image cross-correlation method. Each image was subjected to the background subtraction using white top-hat filter with a radius of 3 pixels and to the spot enhancement by filtering with a Laplacian of Gaussian (LoG) filter in order to increase the signal-to-noise.

RNA Transcripts were detected using local maxima finding method and distinguished from the background noise with an intensity threshold. The minimum intensity threshold was determined plotting dot intensities over threshold values and then choosing the closest value beyond which the number of detected dots showed an evident drop. After a manual cell segmentation using the DAPI and KRT19 images, RNA detected spots were assigned to the cells and were counted in each cell to generate single-cell spatial expression data.

#### Computational methods

##### RNA-seq data analysis

After the pre-processing steps (quality trimming and adapter clipping), the first 3 nt of the second read derived from the G-overhang of the template-switching oligonucleotide were discarded. Paired-end reads with a minimum of 20 bp were aligned to human genome build GRCh38/hg38 using Kallisto (Bray et al., 2016) with default parameters and in the correct orientation (--rf-stranded). The GENCODE version 33 gene annotations was provided as the reference transcriptome. Transcript-level expression was loaded using tximport R package (Soneson et al., 2015) and read counts were normalized using DEseq2’s median of ratios (Love et al., 2014). Normalized counts for protein-coding genes were Log_2_-transformed and used as proxy of expression for downstream analyses. Genome-wide tracks were linearly scaled according the sample size factor.

##### Subtype identification using consensus clustering

The top 25% most variable genes (different iterations from top 30% to top 5% were also tested) in the LMD gene expression dataset were identified using rowVars R function and then used to cluster samples into different subtypes. Consensus clustering was performed using ConsensusClusterPlus R package (Wilkerson and Hayes, 2010) with Pearson’s correlation distance, k-means clustering algorithm and 1’000 re-samplings (100% observation and 100% feature re-samplings). Considering the relative change in area under the cumulative distribution function (CDF) curve for consensus values for different clusters, the number of clusters was determined when there was no substantial increase in consensus.

##### Differential gene expression analysis

Differential gene expression analysis was performed in groupwise fashion across the different subtypes using the DEseq2 R package (Love *et al*., 2014). The patient ID was used as covariate in the model. *p*-values for each tested gene were computed using the Wald test, and Benjamini-Hochberg correction was applied to correct for multiple hypotheses testing. The cutoff for selecting significantly differentially expressed genes was set to adjusted *p*-value (or FDR) ≤ 0.01 and a linear fold change ≥ 2 (in either direction).

##### Unsupervised clustering using k-nearest neighbors

*K*-nearest neighbors (KNN) clustering was used to stratify genes based on their pattern of expression across samples. We first calculated the standardized gene expression considering all the differentially expressed genes. We then performed a Principal Component Analysis (PCA) to identify the top principal components (PCs) that explained the majority of the variance (more than 50%) in the data. We then built a KNN graph of selected genes (*k* = 115) based on the top PCs (3) and performed community detection on this neighborhood graph using the Louvain graph clustering method in igraph R package (https://igraph.org/r/) with default parameters.

##### Functional enrichment analysis and gene signature score

Functional enrichment analyses were performed using EnrichR tool (Kuleshov et al., 2016). We restricted the analysis to Biological Process categories and selected GO terms with enrichment of adjusted *p*-value (FDR) ≤ 0.1. REVIGO (Supek et al., 2011) was used in tiny mode to summarize GO terms and reduce redundancy.

Transcriptional gene signatures reported previously (Bailey *et al*., 2016; Collisson *et al*., 2011; Hwang *et al*., 2022; Moffitt *et al*., 2015) were used to calculate a score for all samples using Seurat’s AddModuleScore function. Default parameters were used except for the number of control features that was set equal to the number of gene signature features. We calculated the standardized gene signature score and plotted as heatmap using the ComplexHeatmap R package.

Enrichment analyses of previously reported transcriptional single cell programs(Hwang *et al*., 2022) in each biotype-specific gene signature (up-regulated DEGs from RNA-seq) were generated using ClusterProfiler’s enricher function (Yu et al., 2012). The adjusted p-value (FDR) was plotted as heatmap using the ComplexHeatmap R package.

##### Supervised learning using Random Forest

A multi-class classifier was implemented to classify all the samples from the RNA-seq. Normalized gene expression matrix (as described above) was preprocessed by centering and scaling samples using the Tidymodels R package (https://github.com/tidymodels). The entire cohort was divided in two equally sized training and testing dataset using a stratified sampling method that preserved the same proportions of samples in each class as in the original data. The classifier was build using a Random Forest (RF) classification method setting a high number of trees (500) and using the k-fold cross validation to avoid overfitting issues. Moreover, the number of features sampled for each tree (mtry) and the minimum number of data points in a node to allow further splitting (min_n) were tuned to optimize the hyperparameters of the model. Given the large amount of features used, the recursive feature elimination (rfe) function in caret R package was used as selection procedure to identify and remove the weakest features. The vip R package (https://github.com/koalaverse/vip) with default parameters was then used to measure the variable Importance as significant impact of features on the model.

To evaluate the model performance, the resulting model was used to predict samples from the test set (**Supplemental Table 5**).

##### Prognostic model

A machine learning-based approach was implemented to develop a prognostic model and to extrapolate prognostic genes starting from the 457 predictor genes of the Random Forest-based model described above using clinical RNA-seq bulk data. Gene expression and clinical data of primary PDAC patients enrolled in the study cohorts, PACA-CA (Pancreatic Cancer Canada) and PACA-AU (Australian Pancreatic Cancer Genome Initiative), were downloaded from the ICGC database while those enrolled in the TCGA study cohort were obtained by the GDC database. Moreover, samples from the PACA-CA cohort that were used in Chan-Seng-Yue et al. (Chan-Seng-Yue *et al*., 2020) were maintained for this analysis (n=118). Normalized gene expression matrix was standardized using the Tidymodels R package (https://github.com/tidymodels). The univariate Cox proportional hazards regression was performed to identify significant genes potentially correlated with the survival of PDAC patients (Cox Univariate, p £ 0.05). Using the genes selected above, the elastic-net (LASSO) regression against the overall survival was used to evaluate simultaneously the survival correlation of multiple features. The model was build using the k-fold cross validation to avoid overfitting issues. After the establishing of the optimal multivariate model using the concordance index (C-index), genes with nonzero model coefficients were further selected to calculate the molecular risk score (combination of gene expression values weighted by the model coefficients) for each patient. A cut-off value was determined as the median risk score. Kaplan-Meier survival curves were generated using the Survminer R package (https://rpkgs.datanovia.com/survminer).

##### GeoMx DSP Data analysis

FASTQ files were trimmed, aligned and deduplicated using GeoMx^®^ NGS Pipeline one Illumina BaseSpace cloud environment as described previously (Merritt et al., 2020). The GeomxTools R package was used to perform pre-processing and to assess the quality of the samples and probes. Samples were retained according to a set of criteria based on the sequencing results (percent of mapping ≥ 80 and minimum number of reads per segment ≥ 1000). Outlier probes were removed on a gene-wise basis to exclude those showing an average count across all segments accounting for less than 10% of the counts of all the probes for the gene. Moreover, a probe was also excluded according to the Grubb’s test in at least 20% of the segments (if the counts of that probe was always higher or lower than the other probes for the same gene). Gene expression was calculated as the geometric mean of the remaining probes and read counts were normalized using DEseq2’s median of ratios (Love *et al*., 2014). Standardized scores of transcriptional gene signatures of three biotypes (up-regulated DEGs from RNA-seq) were calculated for each sample using Seurat’s AddModuleScore function. To cluster all samples in distinct groups, we used the ConsensusClusterPlus R package (Wilkerson and Hayes, 2010) with Pearson’s correlation distance, k-means clustering algorithm and 1000 re-samplings (100% observation and 100% feature resamplings).

##### Gene panel mutational data analysis

After pre-processing (quality trimming and adapter clipping), paired-end reads with a minimum length of 20 bp were aligned to human genome (GRCh38/hg38 build) using bwa mem aligner (Li and Durbin, 2009) with default parameters. Then, Sambamba (Tarasov et al., 2015) was used to remove unmapped reads and PCR duplicates were marked by Picard Tools (http://broadinstitute.github.io/picard). The alignments were subjected to a recalibration of base quality scores using the BaseRecalibrator function (part of GATK4 (McKenna et al., 2010) providing known sites of variation (ExAc data). The resulting BAM files were used to estimate final metrics for each sample using the CollectHsMetrics function of the Picard tools and for the following mutation calling pipeline.

##### Mutation calling pipeline

Mutect2 (part of GATK4) (Cibulskis et al., 2013) was run individually on each tumor sample for the single nucleotide variant calling without a matched normal sample as reference and using the following parameters:

– 1000g_pon.hg38.vcf.gz was downloaded from the Broad Institute Bundle and was used as normal reference (--panel-of-normals option);
– af-only-gnomad.hg38.vcf.gz was used as the source of germline variants with estimated allele frequency (--germline-resource option);
– targeted regions derived from Frige’, et al. (ms in preparation) were specified (-L option).

The LearnReadOrientationModel function of GATK4 was then used to estimate the substitution errors occurring as a result of FFPE artefacts. The resulting orientation bias model was fed into the FilterMutectCalls function of GATK4. The resulting SNVs and INDELs marked as PASS were retained and were further filtered considering a minimum depth of 20 reads and a minimum of 5 reads on the alternate allele.

ANNOVAR (https://annovar.openbioinformatics.org/en/latest) was used to annotate the called variants using the NCBI Refseq release 200 gene annotations. Variants annotated in the gnomAD database with a minor allele frequency (MAF) greater than 0.05 were removed.

To ensure enough coverage in the different samples, only mutational sites covered with at least 20 reads in more than 75% of the samples were retained.

##### Single-cell RNAseq data analysis

Single-cell RNA-seq profiles derived from primary PDACs were mined from Chan-Seng-Yue et al. (Chan-Seng-Yue *et al*., 2020). Count matrices were imported and further processed for downstream analysis, using the Seurat R toolkit (Hao et al., 2021). Low quality cells (defined as showing either >25% reads mapping to mitochondrial genes or <1,000genes expressed) were excluded. Data was then normalized using SCTransform (Hafemeister and Satija, 2019). Principal Component Analysis (PCA) was performed using the normalized expression of the top 2,000 highly variable genes. The top PCs (determined as the number of PCs whose percentage of variance increase among consecutive PCs is less than 0.1%) were then used to build a k-nearest neighbors’ graph (k=20) of single-cell profiles, and to perform community detection using the Leiden clustering method (resolution=0.8), for the identification of distinct cell clusters.

Different cell populations were identified using known cell type-specific gene markers from fibroblasts, immune cells and endocrine/exocrine pancreatic cells (Muraro et al., 2016; Tosti et al., 2021). To confirm the presence of clusters enriched for cancer cells, somatic CNVs were inferred from the single-cell gene expression profiles using the InferCNV R package (https://github.com/broadinstitute/inferCNV). This step was run on all malignant cells using a set of high confidence non-neoplastic cells (identified in the same data) as reference. InferCNV was run with parameters --cutoff 0.1 and --noise_filter 0.2.

##### Single cell classification

Cancer cells from individual patients were scored using the gene signatures of three biotypes we previously identified (up-regulated DEGs from LMD-seq) and for Classical- and Basal-like-related gene signatures (Bailey *et al*., 2016; Collisson *et al*., 2011; Moffitt *et al*., 2015). Seurat’s AddModuleScore function was used as described above to estimate the expression of each gene signature at the single-cell level, properly accounting for the average expression of the genes in each signature, by using a background set of randomly selected genes with comparable expression levels. We then assigned a probability of observing each specific score by chance, given a signature and a single cell, by performing a Permutation test. Briefly, we randomly re-shuffled the gene expression matrix without replacement 100 times, and each time we computed the scores for each signature, for each single-cell. Given a real score, an empirical *p*-value was then associated to it as the number of times a score equal or higher than the real score was observed in the those generated from the permuted data. Each single cell was then classified as a specific biotype, if the *p*-value of the score for the associated signature was <= 0.05.

##### CytoTRACE

CytoTRACE (Gulati *et al*., 2020) was used to predict the developmental potential of single cells ranking them from less differentiated to more differentiated on the basis of their number of expressed genes. After calculating the total number of detected genes for each single cell, CytoTRACE correlates this to the expression of genes to define a gene count signature (GCS), and then it smooths this between 0 (more differentiated) and 1 (less differentiated).

##### CNV estimation

To identify CNV patterns specific to each biotype, CNVs for each single cell were estimated using the inferCNV R package (https://github.com/broadinstitute/inferCNV) and a set of high confidence non-neoplastic cells as reference, as described above. We then computed the mean of CNVs values across all cells of the same biotype, separately for each patient. We then performed hierarchical clustering using Euclidean distance and the Ward.d2 method. The pvclust R package (https://github.com/shimo-lab/pvclust) was then used to assess the cluster stability via bootstrapping (1’000 boostraps). The CNV scoring method (Peng et al., 2019) was used to quantify the overall level of CNVs in each single cell. To this end, CNVs values estimated by inferCNV were first re-standardized to a mean of zero, and then scaled from −1 to 1. In order to account for both deletions and duplications, the sum of squared values was calculated as the overall CNV score for each cell.

##### Motif enrichment analysis

Considering the sequences from −500 bp to +50 bp relative to annotated transcription start sites of differentially expressed genes of the three biotypes (NCBI Refseq release 200 gene annotations), motif enrichment analysis was performed using Pscan (Zambelli et al., 2009) and using a custom set of position-specific weight matrices (PWMs) (Diaferia *et al*., 2016). PWMs were considered significantly overrepresented when showing a *p*-value ≤ 1E-5.

##### Analysis of Regulatory Networks

ARACNe-AP (Lachmann *et al*., 2016) was run with 200 bootstrap iterations for the reconstruction of gene regulatory networks using the gene expression matrix based on laser microdissection (LMD) data. To identify all relationships among regulators and their gene targets, we used a published list of transcription factors (Lambert et al., 2018) that was further manually curated removing well-known chromatin remodelers and mitochondrial transcriptional regulators (final number of TFs was 1590). ARACNe networks were imported and used by the VIPER package (Alvarez *et al*., 2016) for the identification of possible master regulators. We compared each biotype against each other using the msviper algorithm of the VIPER package and the gene signatures previously identified. To infer the enrichment of the top differentially expressed transcription factors, we measured the activity of a regulator based on the enrichment of its target genes in each biotype. Representative differentially expressed transcription factors and their gene targets were displayed as networks using cytoscape (v.3.8.2) (Shannon et al., 2003).

##### Multiplexed RNA smFISH Data Analysis

Single-cell expression matrix and spatial coordinates were imported into the Giotto (Dries et al., 2021) R package and were further processed for downstream analysis. Low quality cells (≤5 genes detected per cell and at least 1 molecule as expression threshold) were excluded. The data were normalized per gene and per cell using osmFISH (Codeluppi et al., 2018). Z-scored gene expression was used for the Principal Component Analysis (PCA) and for the finding of the top PCs (20). Cell cluster identification was performed by k-nearest neighbors graph (k=30) and community detection using the Leiden graph clustering method (res=0.7). UMAP manifold was used for visualization of the cluster groups. Markers from each cluster were identified according to the findMarkers_one_vs_all function, intersected with the genes up-regulated in LMD-seq and then used to assign cell cluster to the three biotypes. To infer tumor areas based on the spatial coordinates of the identified cell types, single cells belonging to the identified cluster groups were mapped back to their original histological space. Moreover, gene expression of selected genes was plotted by spatFeatPlot2D function.

##### Statistical analysis

Unless indicated otherwise, statistical analyses were performed in the statistical computing environment using R v4.

## Supplementary Figures

**Supplementary Figure 1.**
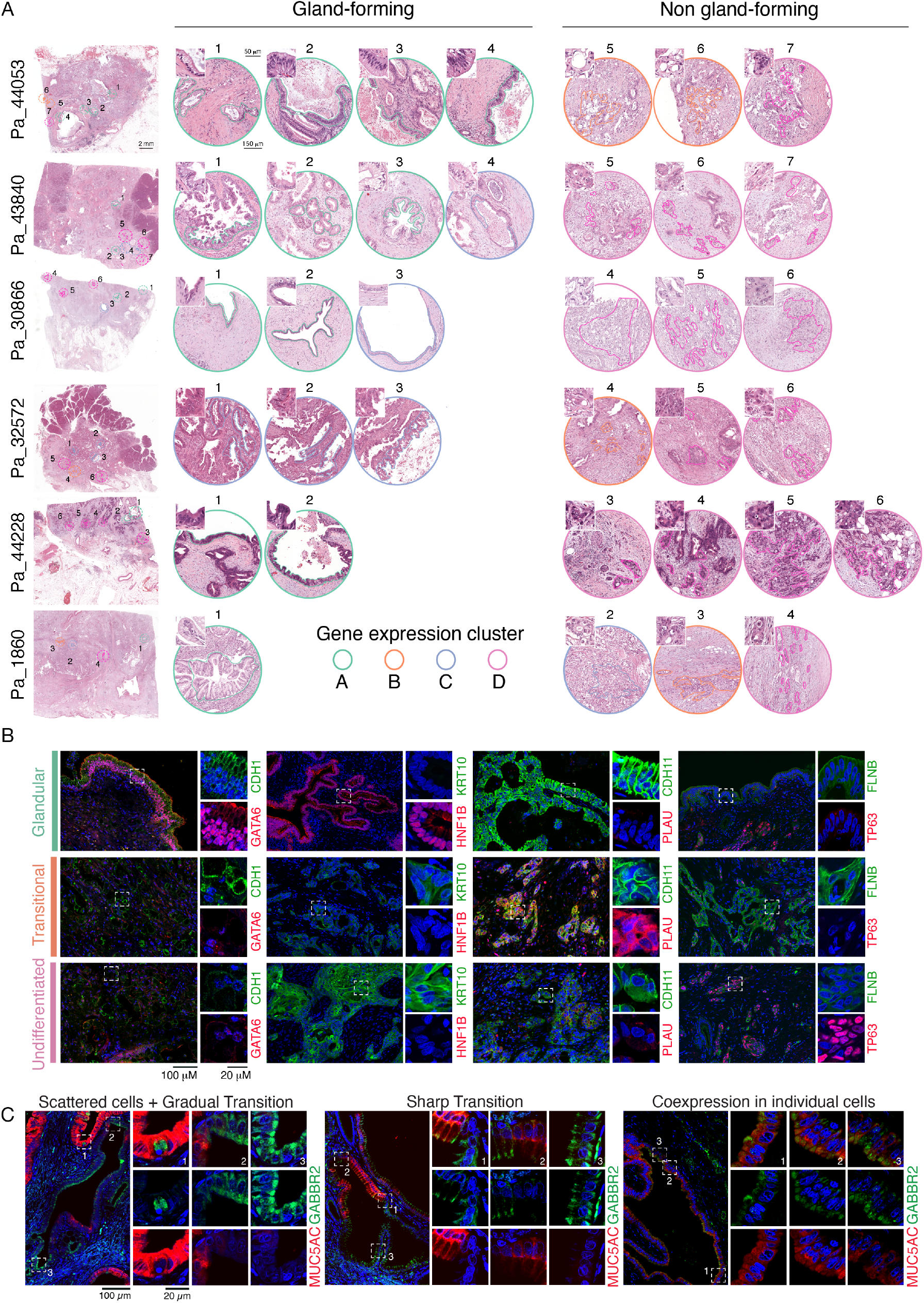
Representative PDAC cases used for LMD-seq. **A**) H&E-matched images of the microdissected areas from 6 representative patients, divided into gland-forming and non gland-forming areas. Border lines indicate the microdissected areas, while the colors match the assigned biotype (A = green, B = orange, C = light blue, D = pink). Scale bars: overview images 2mm, round images 150 μm, insets 50 μm. **B**) Immunofluorescence analysis of protein markers representative of the different biotypes. Large images show the merged channels with DAPI-counterstained nuclei (blue). Scale bar: 100μm. White boxes indicate the area shown at higher magnification on the right, with red and green channels split in two images. Scale bar: 20 μm. **C**) Representative immunofluorescence images of microdissected areas matching cluster C (hybrid). Different tumor areas were immunostained and probed for GABBR2 (green) and MUC5AC (red). Large images show the merged channels with DAPI-counterstained nuclei (blue). Scale bar: 100μm. White boxes indicate the area showed at higher resolution on the right. Scale bar: 20 μm.

**Supplementary Figure 2.**
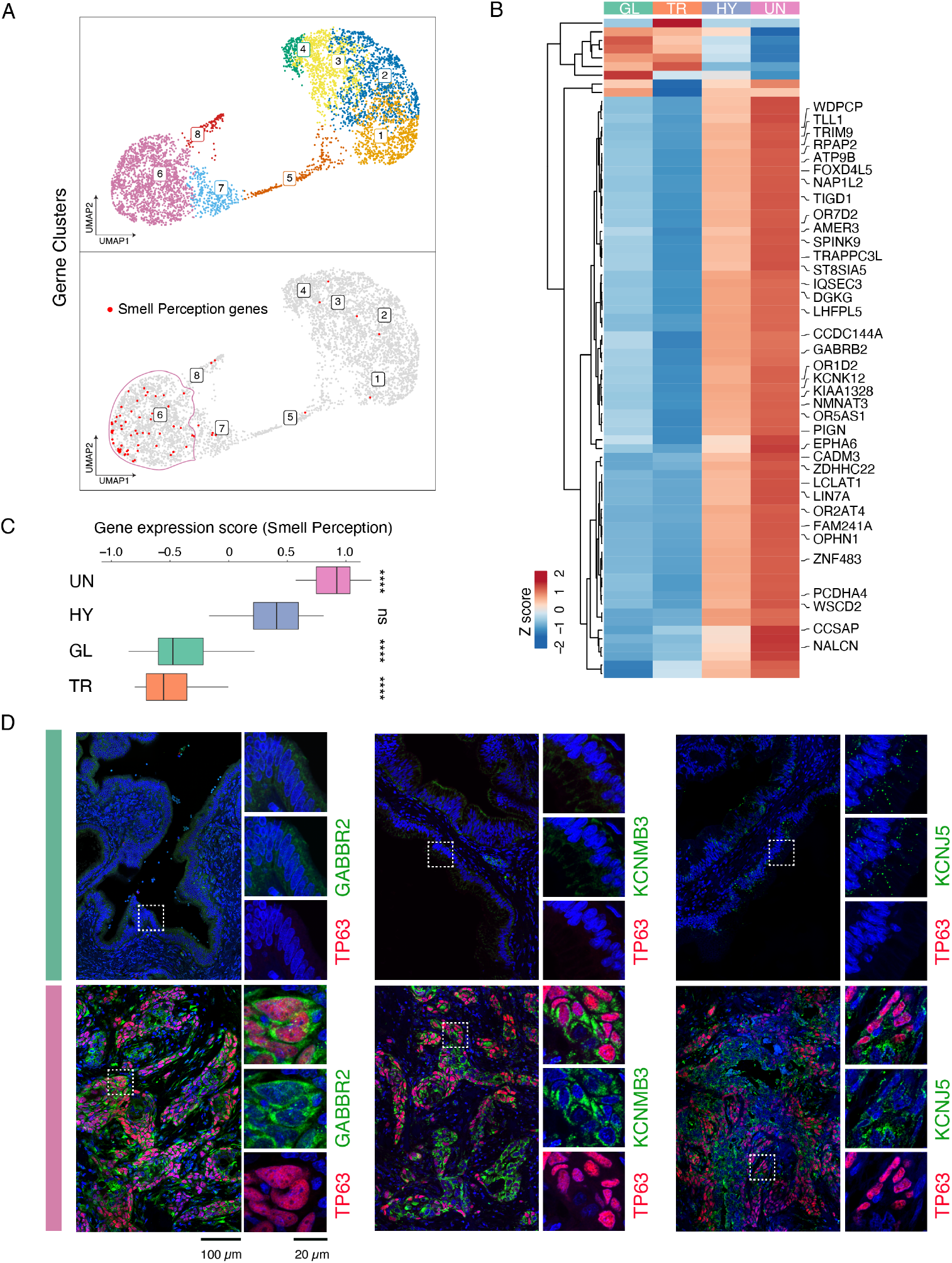
Selective expression of the smell perception gene expression program in the undifferentiated biotype. **A**) UMAPs showing the separation of the differentially expressed genes belonging to their specific gene clusters (from 1 to 8 as shown in **Figure 1**) (top) and the enrichment of 76 differentially expressed genes involved in the smell perception gene expression program annotated in the Human Protein Atlas (HPA; https://www.proteinatlas.org/humanproteome/tissue/expression+cluster#cluster6) (bottom). **B**) Heatmap showing differentially expressed smell perception genes in PDAC biotypes. Genes with evidence at protein level in Uniprot or HPA are shown. **C**) Box plot showing the gene expression scores of the Smell Perception gene expression program calculated for each sample and stratified by biotype. Kruskal-Wallis P-value and two-sided Wilcoxon rank-sum tests (group versus rest) are shown. Biotypes are ordered from top to bottom based on decreasing gene expression score level. **D**) IF staining of selected voltage-gated ion channels enriched in the undifferentiated biotype (lower row, pink bar) compared to the glandular biotype (upper row, green bar). Large immunofluorescence images showed the merged colors with DAPI-counterstained nuclei (blue). Scale bar: 100μm. White boxes indicate the area showed at higher resolution on the right, with red and green channels split in two images. Scale bar: 20 μm.

**Supplementary Figure 3.**
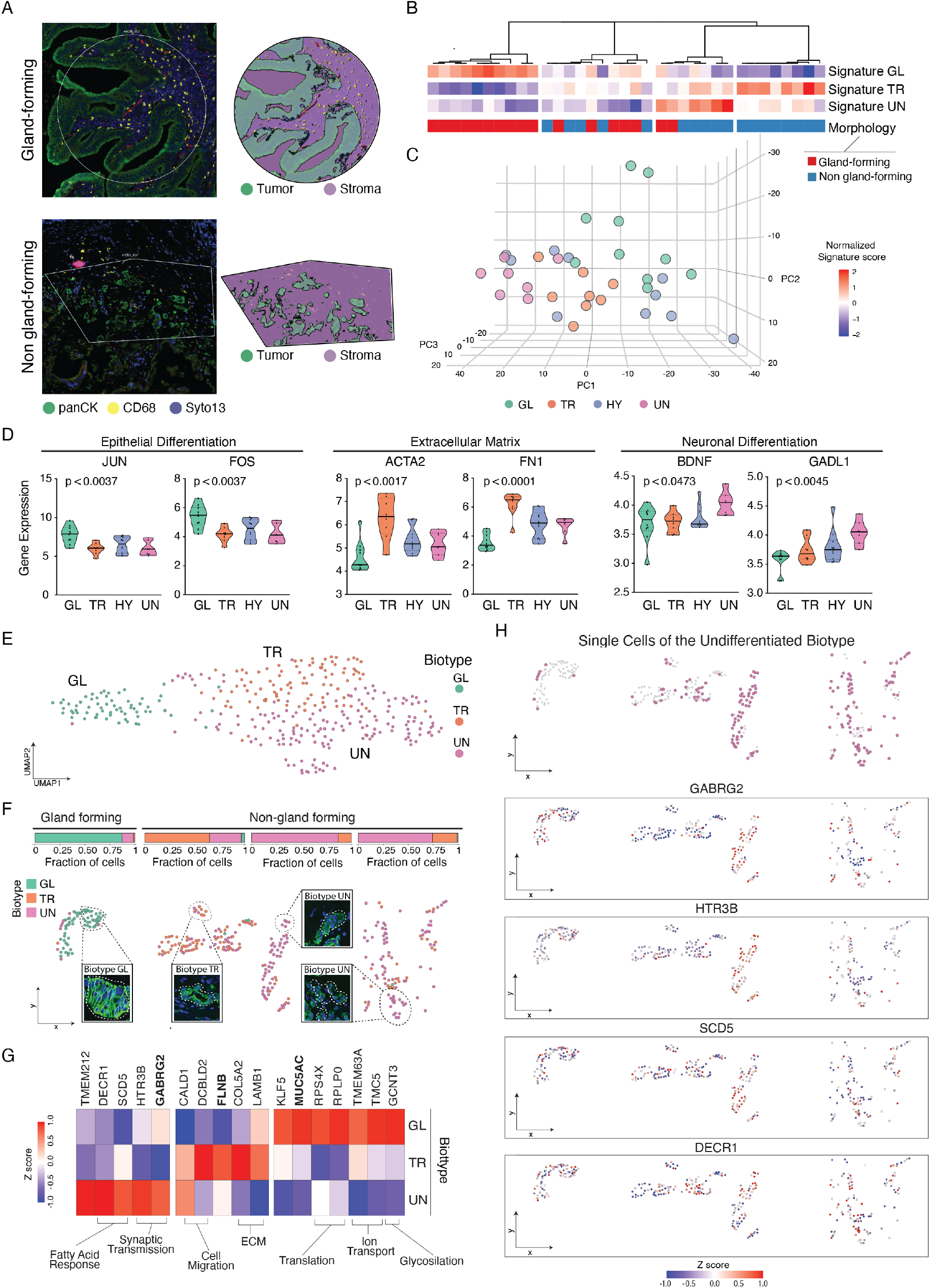
Validation of the PDAC morpho-biotypes with orthogonal methods. **A**) Left, immunofluorescence images with selected regions of interest (ROIs) of gland-forming and non gland-forming tumor areas labelled with anti-pan-CK (green), anti-CD68 (yellow) and Syto13 as nuclear marker (blue). Right, segmentation masks obtained by GeoMx DSP used to enrich for epithelial tumor (pan-CK-positive) cells and stromal (pan-CK- and CD68-negative) cells. **B**) Unsupervised clustering of 35 samples from 8 PDAC patients captured by the GeoMx DSP. Samples were first scored for the three gene signatures derived from the up-regulated genes in the three biotypes (signature GL, signature TR and signature UN) and then clustered using ConsensusClusterPlus (Wilkerson and Hayes, 2010) with Pearson’s correlation distance, k-means clustering algorithm and 1000 iterations. Morphology (gland forming and non-gland forming) is shown for each sample. **C**) 3D-PCA plots of the 35 samples showing the separation of four identified clusters that were assigned to each biotype according to the gene signature expression (see **B**). **D**) Violin plots showing the expression levels in different biotypes of two representative genes in each of the three following functional categories: epithelial differentiation (left), extracellular matrix (middle) and neuronal differentiation (right). Gene expression represents the log_2_(normalized read counts scaled by cDNA length + 1). Kruskal-Wallis P-values are indicated for each gene. **E**) UMAP plot visualization of multiplexed RNA single molecule FISH in four distinct heterogenous tumor areas of a PDAC patient (n=331 single cells). Three single-cell clusters were identified and assigned to the three morpho-biotypes according to marker gene expression (see **G**). **F**) Spatial organization of the three morpho-biotypes in the four tumor areas. Single cells were first mapped back to the tissue sections and then labeled according to cluster assignment. A highlight of the representative morphologies of some single cells belonging to the three biotypes is shown. The fraction of single cells assigned to the three biotypes in each tumor area is shown above the spatial organization plots. **G**) Heatmap of differentially expressed marker genes identified in each morpho-biotype and grouped by their functional annotations. Genes also validated at the protein level are shown in bold. **H**) Highlight of the spatial organization of single cells belonging to the UN biotype (above) and the spatial expression of four distinct markers (bottom) representing the following functional categories: synaptic transmission (GABRG2 and HTR3B) and fatty acid response (SCD5 and DECR1).

**Supplementary Figure 4.**
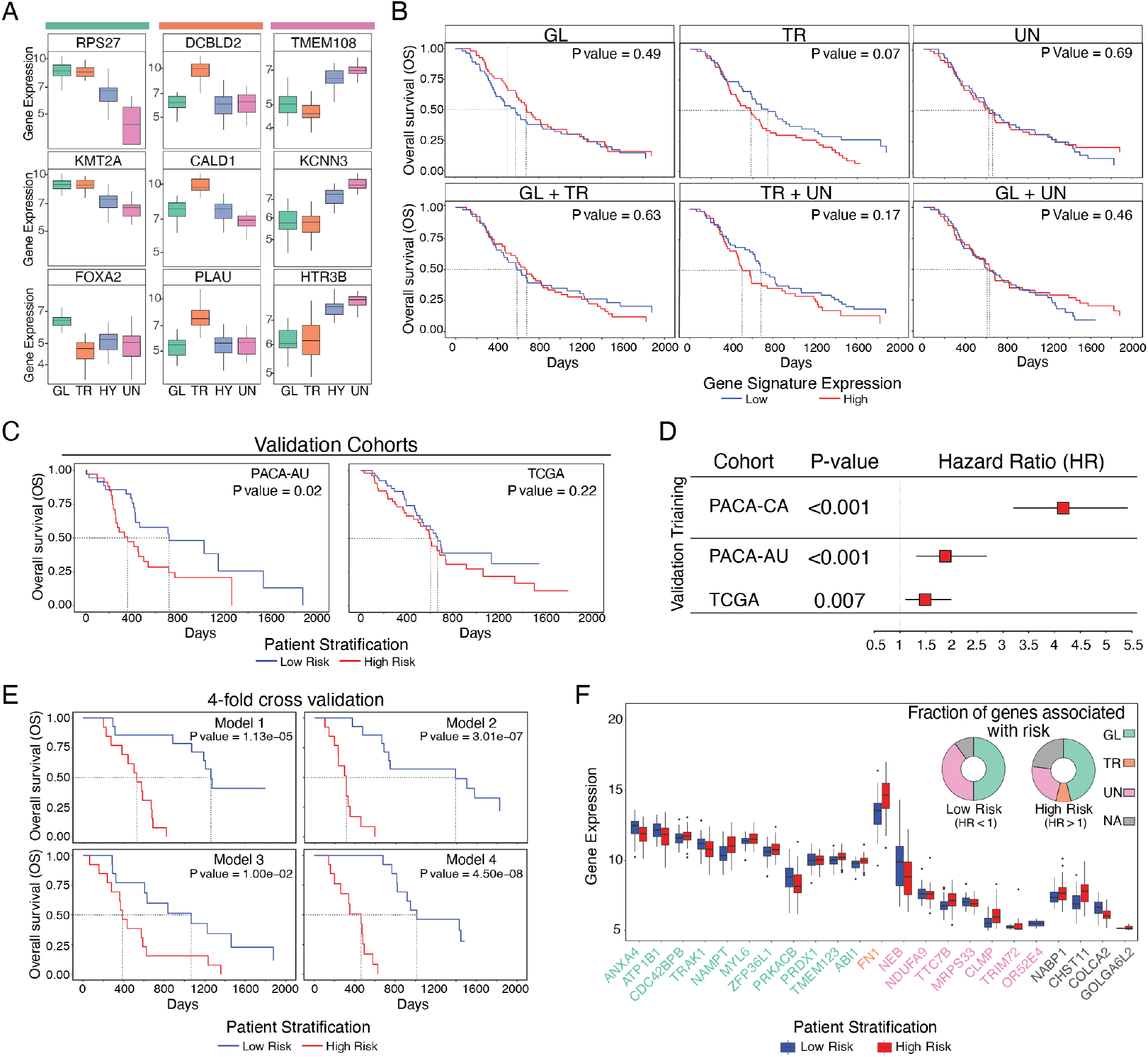
Validation of the prognostic relevance of the PDAC morpho-biotypes. **A**) Box plots showing the expression levels of representative predictor genes in each biotype. The Glandular biotype corresponds to cluster A (green), the transitional biotype to cluster B (orange) and the undifferentiated biotype to cluster D (pink). Gene expression represents the log2(normalized read counts scaled by cDNA length + 1). **B**) Kaplan-Meier plots of overall patient survival stratified by the expression of single or combined biotypes gene signatures (PACA-CA dataset, n=110). Log rank P-value is shown. Dashed lines indicate the median survival for each biotype. GL: Glandular, TR: Transitional, UN: Undifferentiated. **C**) Kaplan-Meier plots of overall patient survival for high-risk and low-risk groups in two independent validation cohorts (PACA-AU dataset, n=70; TCGA dataset, n=90). Log rank P-value is shown. Dashed lines indicate the median of survival for each group. **D**) Univariate Cox regression analysis of the gene signature risk score on the training (PACA-CA dataset, n=110) and validation cohorts (PACA-AU dataset, n=70; TCGA dataset, n=111). P-value and Hazard Ratio (HR) with 95% confidence interval are shown. **E**) Kaplan-Meier plots of overall patient survival for high-risk and low-risk groups (PACA-CA dataset, n=110) in four distinct prognostic models generated by dividing the cohort into four different testing sets (n=27 for each model). **F**) Expression level of the 23 prognostic genes in high-risk and low-risk patients. The genes are ranked by morpho-biotype (GL: Glandular, TR: Transitional, UN: Undifferentiated) and expression level. The donut charts show the fraction of genes belonging to the three biotypes associated with the risk (Low risk: HR < 1; High risk: HR > 1). HR: Hazard Ratio.

**Supplementary Figure 5.**
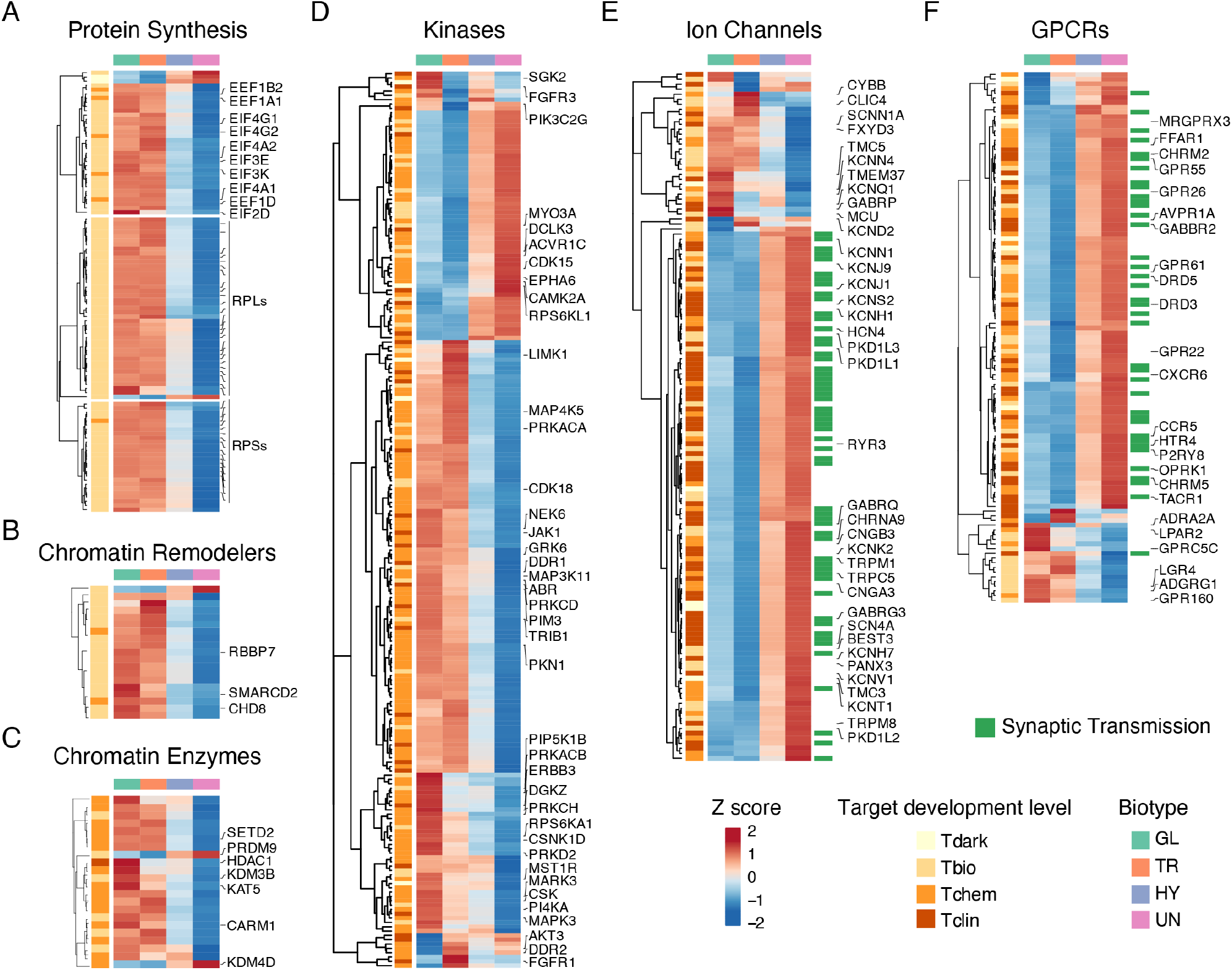
Target development level of genes differentially expressed in PDAC biotypes. **A-F**) A knowledge-based classification system was used (https://pharos.nih.gov) (Oprea et al., 2018) that divides gene products into four different target development levels, ranging from T_dark_ (gene products with no biological annotation) to T_clin_ (gene products targeted by at least one clinically approved drug), with two intermediate categories represented by T_bio_ (gene products associated with extensive biological information) and T_chem_ (proteins bound and affected by a chemical molecule). Shown are hierarchically clustered heatmaps (Euclidean distance and complete linkage method) of differentially expressed genes grouped into six functional categories: protein synthesis (**A**), chromatin remodelers (**B**), chromatin enzymes (**C**), kinases (**D**), ion channels (**E**), GPCRs (**F**). Highly differentially expressed genes (fold change ≥ 3) for each category are shown. For ion channels and GPCRs, genes related to synaptic transmission are flagged.

**Supplementary Figure 6.**
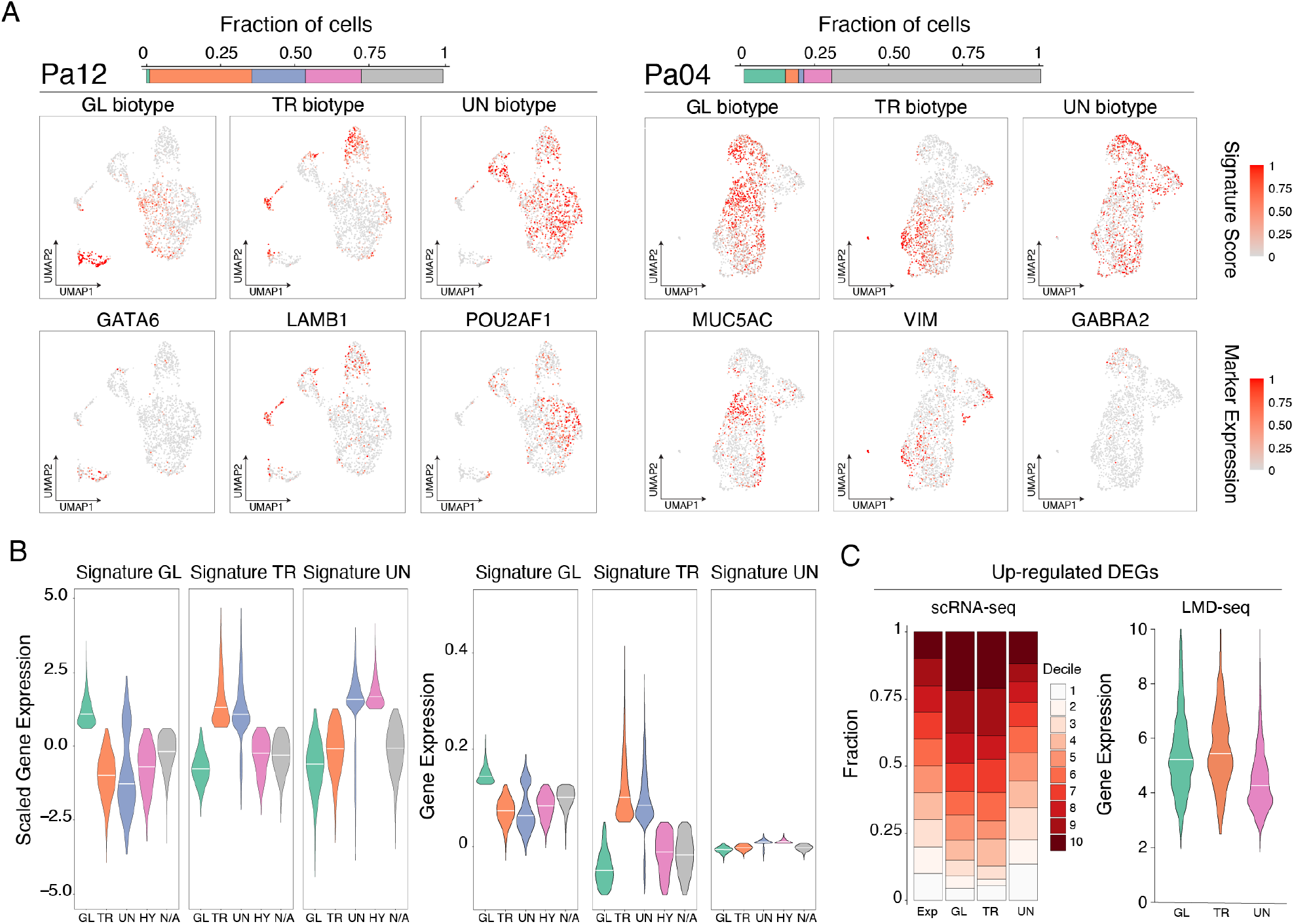
Enrichment of gene expression signatures in scRNA-seq data. **A**) UMAPs of malignant cells from 2 selected patients labeled by the gene expression score obtained for each biotype-specific gene signature (top) (GL: Glandular, TR: Transitional, UN: Undifferentiated) or by expression of selected genes characteristic for each biotype (bottom). **B**) Violin plots showing the scaled (left) or normalized (right) expression of the gene signatures across the classified cells (**Figure 3C, D**). **C**) Distribution of gene expression levels of up-regulated DEGs for each biotype defined in LMD data using the scRNAseq expression matrix (left) or the LMD expression matrix (right).

## Supplementary Table Legends

**Suppl. Table 1**. *Patients included in the study*.

**Suppl. Table 2**. *Sequencing summary*. Summary of LMD RNA-seq in laser microdissected PDAC samples. Sample and sequencing annotations (including sequencing depth, unique mapped reads and number of expressed genes) are shown.

**Suppl. Table 3**. *Differentially expressed genes in PDAC biotypes*. Gene expression analysis in laser microdissected PDAC samples. Biotype assignments based on the top variable genes and the differentially expressed genes (DEGs) are shown.

**Suppl. Table 4**. *Functional categories associated with PDAC biotypes*. Functional enrichment analysis was carried out using the GO-biological processes in the MSigDB database. The table includes GO terms summarized by their semantic similarity and the complete enrichment analysis for each selected cluster of differentially expressed genes.

**Suppl. Table 5**. *A random forest classifier of PDAC biotypes: selected features and their importance*. Variable Importance and statistics on the testing set are reported.

**Suppl. Tables 6**. *Target development level of differentially expressed genes characteristic of different PDAC biotypes*. Differential expressed genes were grouped into categories related to therapeutic vulnerabilities of the encoded proteins. Target development levels based on the Pharos database annotations and gene expression levels in different biotypes are shown for each gene.

**Suppl. Table 7**. *Mutational profiles associated with PDAC biotypes*. Mutational analysis in laser microdissected PDAC samples was carried out using a custom amplicon-based panel of 467 genes. Sequencing summary with sample and sequencing annotations (including sequencing depth, coverage and number of identified mutational sites), somatic variants annotated by ANNOVAR and groupwise Fisher’s exact tests are shown.

**Suppl. Table 8**. *Transcriptional regulatory networks enforcing distinct PDAC biotypes*. The table provides in the different sheets the following information: a) the list of all differentially expressed transcription factor genes; b) the position weight matrixes (PWM) used in the motif enrichment analyses and c) the master regulators derived by the VIPER analysis in the different biotypes. For each PWM the source (database or publication) is indicated while for each master regulator the gene expression levels and log2FC are shown.

